# Metabolic control of YAP via the acto-myosin system during liver regeneration

**DOI:** 10.1101/617878

**Authors:** Kirstin Meyer, Hernan Morales-Navarrete, Sarah Seifert, Michaela Wilsch-Braeuninger, Uta Dahmen, Elly M. Tanaka, Lutz Brusch, Yannis Kalaidzidis, Marino Zerial

**Affiliations:** Max Planck Institute of Molecular Cell Biology and Genetics, Dresden, Saxony 01307, Germany; Experimental Transplantation Surgery, Department of General, Visceral and Vascular Surgery, Jena University Hospital, 07747 Jena, Germany; Research Institute of Molecular Pathology, Vienna Biocenter, 1030, Vienna, Austria; Faculty of Bioengineering and Bioinformatics, Moscow State University, 119991 Moscow, Russia; Center for Information Services and High Performance Computing, Technische Universität Dresden, 01062 Dresden, Germany

## Abstract

The mechanisms of organ size control remain poorly understood. A key question is how cells collectively sense the overall status of a tissue. We addressed this problem focusing on mouse liver regeneration, which is controlled by Hippo signalling. Using digital tissue reconstruction and quantitative image analysis, we found that the apical surface of hepatocytes forming the bile canalicular network expands concomitant with an increase of F-actin and phospho-Myosin, to compensate an overload of bile acids. Interestingly, these changes are sensed by the Hippo transcriptional co-activator YAP, which localizes to the apical F-actin-rich region and translocates to the nucleus in dependence of the acto-myosin system. This mechanism tolerates moderate bile acid fluctuations under tissue homeostasis, but activates YAP in response to sustained bile acid overload. Using an integrated biophysical-biochemical model of bile pressure and Hippo signalling, we explained this behaviour by the existence of a mechano-sensory mechanism that activates YAP in a switch-like manner. We propose that the apical surface of hepatocytes acts as a self-regulatory mechano-sensory system that responds to critical levels of bile acids as readout of tissue status.

## Introduction

Organ size control is a fundamental aspect of morphogenesis. It requires a control system acting throughout scales of organization to coordinate local cell behavior with global tissue properties. Structural, biochemical and mechanical properties of tissues regulate cell proliferation and differentiation (1,2), cell geometry (3) and the growth and organization of cells into tissues (4). Mechanical properties of tissues emerge from the interaction of cells and their environment. These include local cell-cell and cell-matrix interactions as well as systemic factors such as blood pressure, fluid shear stress and muscle tension. Cells respond to biochemical or mechanical alterations through the activation of signaling pathways and downstream effectors, of which one of the most important is the actin cytoskeleton. It is able to generate forces via the acto-myosin system and controls cell behavior through signaling cascades, such as the Hippo and SRF pathways. Current concepts on the reciprocal regulation of tissue structure and cell behavior are primarily derived from studies *in vitro* and *ex vivo* (5,6). However, the responses of cells to the structural and mechanical properties within tissues are largely unexplored.

Liver regeneration provides an excellent example of organ size control. Liver mass scales with body size, constituting about 5% of the body weight in rodents (7) and has the capacity to regenerate the original mass (8). The liver consists of functional units or *lobuli*. Each lobule contains two opposing fluid networks, the sinusoidal endothelial and bile canaliculi (BC) networks that transport blood and bile between portal and central vein (PV-CV), respectively. The hepatocytes are polarized cells at the interface of both networks. They take up metabolites from the blood via their basal plasma membranes and secrete waste products and bile via their apical membranes, which collectively, form a continuous network that drains into bile ducts.

Several signalling pathways are required for liver regeneration (9,10) of which the Hippo pathway is a key regulator of liver size (11,12). The Hippo pathway is activated during liver regeneration (13) to promote hepatocyte proliferation primarily via the co-transcriptional activator yes-associated protein (YAP). It is a mechano-sensor for signals from the actin cytoskeleton, cell polarity complexes and cell-cell or cell-matrix junctions (14–16) and is also responsive to high concentrations of bile acids (BA) (17). Despite these molecular insights, how cells sense the overall tissue status and size remains an open question. In particular, it is unclear how cells integrate metabolic (BA levels) and/or mechanical alterations, like changes in blood and bile pressure, to control liver size during homeostasis and regeneration. Here, we addressed these questions by applying high-resolution microscopy and quantitative 3D image analysis (18,19) to explore tissue and cellular alterations during liver regeneration.

## Results

### Alterations of the BC network during liver regeneration

Tissue sections of liver from different time points after partial hepatectomy (PH) (0.8-5d post PH) were stained for the apical marker CD13 by immunofluorescence (IF), imaged at high resolution by confocal microscopy and the 3D BC network reconstructed form IF image stacks. The analysis revealed remarkable changes of BC network topology and geometry. The BC network dilated and branched as early as 0.8d post PH (Fig.1a) throughout the entire CV-PV axis and reversed to normal at ∼3-5d post PH. Spatial quantification revealed an increase of BC diameter by up to 27% (zone 11) and on average by 21% throughout the entire CV-PV axis at 1.5d after PH as compared to the untreated liver (1.96-2.27µm in untreated liver; 2.40-2.61µm at 1.5d post PH,; Fig.1b). At 5d post PH, the BC diameter was still increased, but only by 11% on average (BC diameter at 5d post PH, 2.14-2.51µm). In addition, the BC lacked the typical smooth and regular appearance at this resolution and acquired a rough surface texture (Fig.1a, compare BC in PV area of untreated vs. 1.5d post PH). This could reflect alterations of the acto-myosin system, which mediates apical contractility and regulates BC geometry and bile flow (18,27). Interestingly, the observed structural changes preceded the reported temporal profile of hepatocyte proliferation, which starts at 1.5d post PH and peaks at 2d (28). The structural alterations of the BC network may just be an epiphenomenon or play an active role in liver regeneration. Therefore, we set to analyze the nature and cause of BC network expansion.

**Figure 1.**
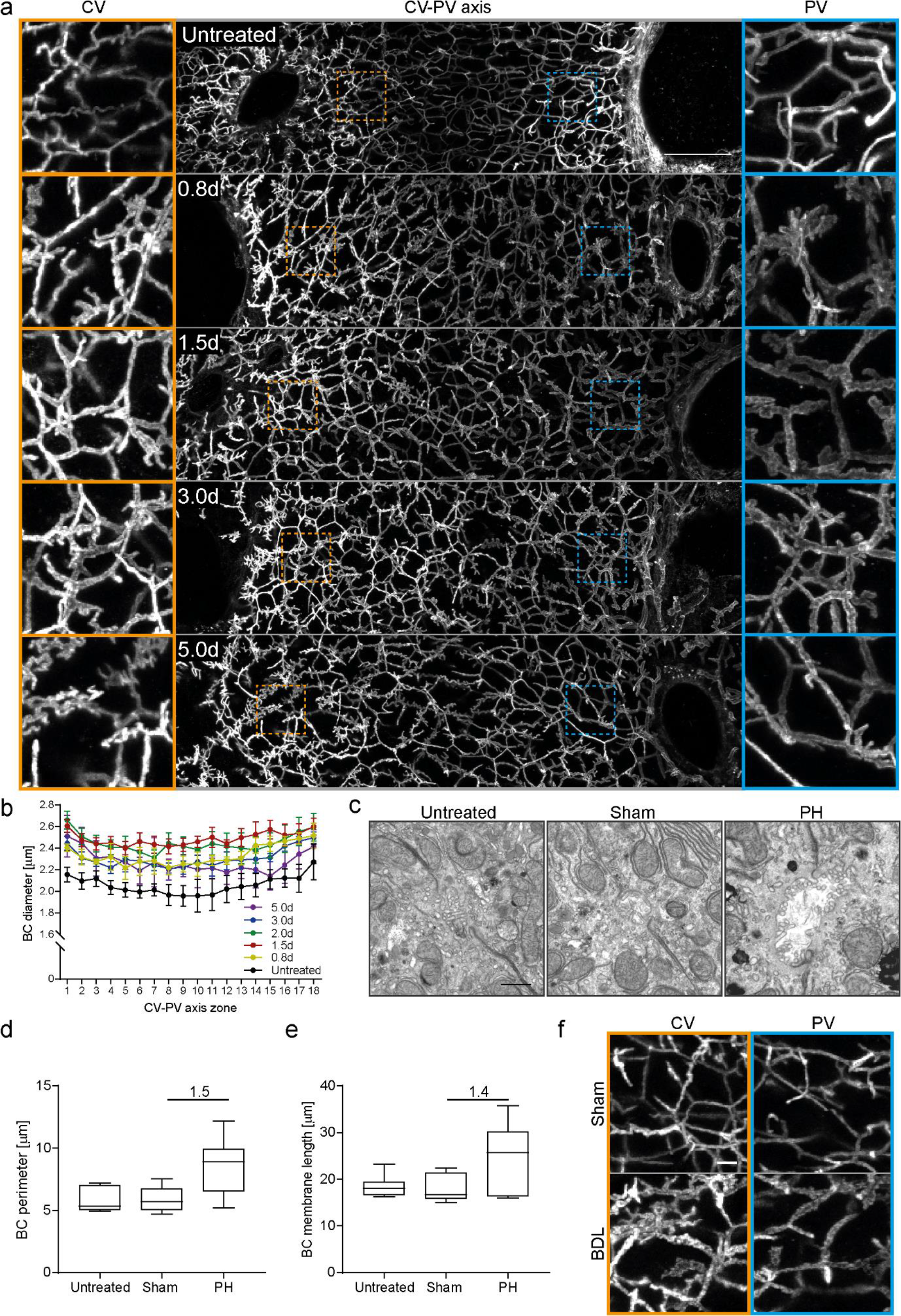
The BC network transiently expands during liver regeneration. **a)** Fluorescence staining for the apical marker CD13 on liver tissue sections from untreated mice or animals at indicated time points post PH. Shown are maximum projections of 50 µm z-stacks covering an entire CV-PV axis (CV, left; PV, right). Indicated regions of the CV (orange) and PV (blue) areas are shown as magnifications on the left and right, respectively. **b)** Quantification of BC diameter within 18 zones along the CV-PV axis (zone 1, peri-central; zone 18, peri-portal) in livers from untreated animals or mice at indicated timepoints post PH. The diameter was measured from 3D BC network reconstructions of IF image stacks of CD13 as shown in (a). The zones directly adjacent to the CV and PV were excluded from the analysis (∼ 1 cell layer). Mean ± s.e.m, n=3-6 mice per timepoint. BC diameter of untreated mice vs. 0.8 d, 1.5 d, 2.0 d, 3.0 d and 5.0 d post PH, p < 0.0001. **c)** EM images of BC on liver tissue sections of untreated mice (left) and 1.8 d post sham OP (middle) or PH (right). **d, e)** Quantification of BC perimeter (d) and total BC membrane length (e) from EM images as representatively shown in (c). Box-whisker plot with median, 25-75 quartiles and minimum/maximum error bars, n=5-6 mice per condition. In (d), BC perimeter of untreated vs. sham condition, p > 0.05 (n.s.); Untreated vs. PH condition, p < 0.01. In (e), BC membrane length of untreated vs. sham condition, p > 0.05 (n.s.); Untreated vs. PH condition, p < 0.05. **f)** IF stainings of the apical marker CD13 on liver tissue sections from livers at 1 d post sham OP or BDL. Shown are maximum projections of 50 µm z-stacks in the CV (left, orange) and PV (right, blue) region. Images in (a) and (f) are background-subtracted. Scale bars, 50 µm (a), 1 µm (c), 5 µm (f).

### Expansion of the apical surface of hepatocytes during liver regeneration

The observed BC diameter increase could be due to a bona fide expansion of the apical surface area of hepatocytes or be only apparent, e.g. due to decreased contractility of the actomyosin cortex and/or flattening of microvilli. To distinguish between these possibilities, we quantified BC membrane length and perimeter by electron microscopy (EM) at 1.8d post PH or control operated (sham) and untreated mice (Fig.1c-e). The apical membrane length was determined by segmentation of the BC, whereas the BC perimeter was estimated by calculating the minimal enclosing ellipse of these segmentations (Fig.S1). The EM analysis (Fig.1c) confirmed the BC expansion (Fig.1a). Both BC perimeter and total membrane length increased to a similar extent, 1.5 and 1.4-fold, respectively (Fig.1d, e). Thus, the primary cause of network dilation is an increase in total apical membrane of hepatocytes.

Liver resection induces a transient overload of BA (29). The observed BC network expansion could be a compensatory response to increased apical secretion and changes of biliary fluid dynamics. To test if BA overload can replicate the BC network expansion, we examined the BC network structure in cholestatic livers, using the model of bile duct ligation (BDL). In this model, the common bile duct is ligated, resulting in BA overload of the organ. After BDL, the BC network in the liver was branched and expanded (Fig.1f), strikingly resembling regenerating liver. As in regeneration, the expansion occurred already at 1d post-BDL throughout the entire CV-PV axis, preceding hepatocyte proliferation (30). Based on these results, we hypothesize that the expansion of the BC network could be part of a mechano-sensory system that responds to increased BA levels to induce liver growth during regeneration.

### Expansion of the apical surface of hepatocytes is associated with increased acto-myosin levels

BC possess a dense sub-apical contractile actin mesh via the acto-myosin system (18,27) which contributes to BC geometry and bile flow (18). To test whether the actin cytoskeleton is modified along with the changes in BC during regeneration, we quantified the levels of apical F-actin (phalloidin, Fig.2a, Fig.S2a) and phospho-Myosin light chain (pMLC, Fig.2c, Fig.S2b) at the apical area of hepatocytes at different time points after PH or sham operation. We had to take into consideration the effects of the sham operation itself on the actin cytoskeleton: The apical F-actin density progressively decreased by up to 37±9% (mean±s.e.m.) compared to untreated mice (Fig.2b), presumably due to the laparotomy and/or anesthesia/analgesia. However, upon PH, the F-actin intensity was higher than in sham operation and fluctuated at baseline levels as compared to untreated mice, with a maximum increase of 17±10% (mean±s.e.m) at 1.5d (Fig.2b). In comparison to F-actin, apical pMLC intensity increased more dramatically during regeneration (Fig.2c, d). Whereas in sham-operated mice, apical pMLC levels remained similar to baseline levels, within a maximum increase of 30±9% (mean±s.e.m.) and decrease of 13±11% (Fig.2d, Fig.S2b), upon PH they raised early and remained elevated by about 70% until ∼2.8d post PH as compared to untreated mice (Fig.2d). Overall, the expansion of the BC network during regeneration is associated with a concomitant increase in apical F-actin and pMLC levels, i.e. acto-myosin contractility.

**Figure 2.**
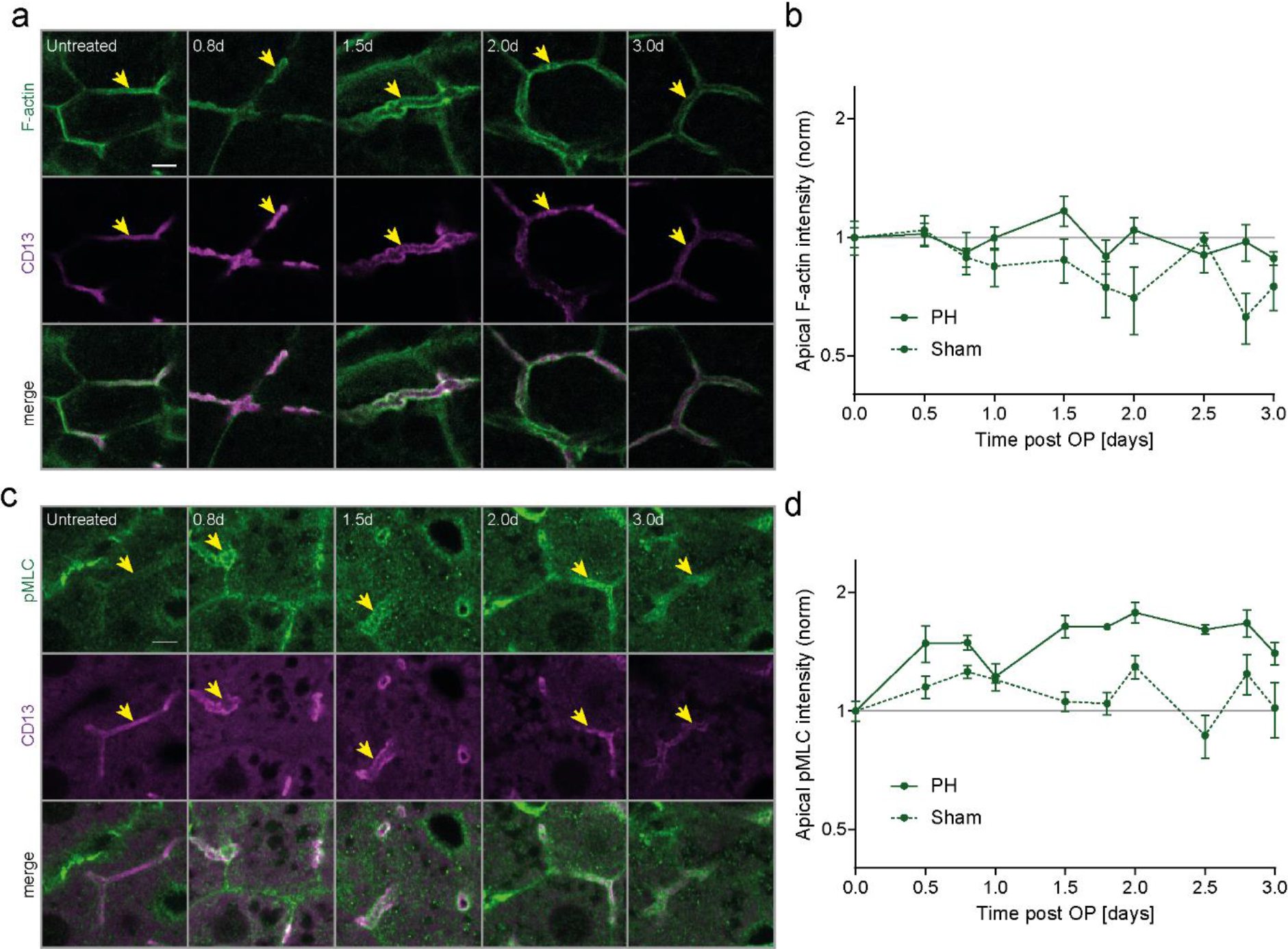
BC network dilation is accompanied by increased apical myosin levels during regeneration. **a, c)** Fluorescence stainings for F-actin (a) or pMLC (c) and the apical marker CD13 in the PV area on liver tissue sections from untreated mice or animals at indicated time points post PH. Arrows indicate BC. **b, d)** Quantification of apical F-actin (b) and pMLC (d) intensity from images as representatively shown in (a,c) as well as Fig.S2a,b, at indicated time points post PH (solid line) or sham OP (dashed line). Data is normalized to untreated animals (timepoint 0). Mean ± s.e.m, n=3-5 (b) and n=2-5 (d) mice per timepoint. Apical F-actin of sham vs. PH time course, p < 0.05; Apical pMLC of sham vs. PH time course, p < 0.005. Scale bar, 5 µm (a,c).

### BC network expansion correlates with YAP activation and proliferation during regeneration

The actin cytoskeleton converts mechanical forces into biochemical signals. Since the Hippo pathway is a prominent mechano-sensor downstream of the actin cytoskeleton and can be activated by BA (17), we hypothesize that it could sense BA indirectly, through their effects on the apical acto-myosin system of hepatocytes, and drive their proliferation. To test whether YAP can sense BC network alterations, we examined its spatio-temporal dynamics using a specific antibody, validated by loss of signal upon YAP KO (Fig.S3). We correlated YAP nuclear localization, as readout for its activity (31,32), with hepatocyte proliferation, detected by the cell cycle marker PCNA. To account for previously reported spatial heterogeneities of proliferation within the liver lobule (33), we imaged the entire CV-PV axis at 17 time points (0.5-7d) during regeneration (Fig.3a, b) and upon sham-surgery (Fig.S4a, b). Consistent with earlier reports (34), YAP was highly expressed in cholangiocytes but at low levels in hepatocytes in the livers of untreated mice (Fig.3a). In hepatocytes, the signal was primarily cytosolic, as expected for non-proliferating cells (35). During regeneration, however, YAP levels increased in hepatocytes as compared to untreated and sham-operated livers as determined by both IF (Fig.3a, Fig.S4a) and Western blot analysis (Fig.S4c). The major fraction of YAP was still cytoplasmic but, in addition, it was also detectable in nuclei of proliferating hepatocytes (Fig.3b, Fig.S4b). Quantification of nuclear YAP and PCNA levels showed that nuclear YAP was specific to regenerating livers compared to sham-operated or untreated mice (Fig.3c). It started as an immediate response to liver resection, as early as 0.5d post PH, ceased with the proliferative wave at about 3-4d (Fig.3c), and occurred throughout the entire CV-PV axis (Fig.3d). These spatio-temporal dynamics remarkably correlate with those of BC network expansion (see Fig.1b). Interestingly, we found that YAP was enriched at the apical plasma membrane of hepatocytes in regenerating liver (Fig.3e). The apical localization was detectable throughout the CV-PV axis, both in PCNA-positive and negative hepatocytes, and was specific to regenerating liver as compared to untreated and sham operated mice (Fig.S5a). Also, YAP displayed a particulate staining, suggesting that it may be spatially concentrated, e.g. on the actin cytoskeleton and/or organelles underneath the apical membrane (Fig.3e). This suggests that the apical localization and activation of YAP may be linked to the alterations of the BC network.

**Figure 3.**
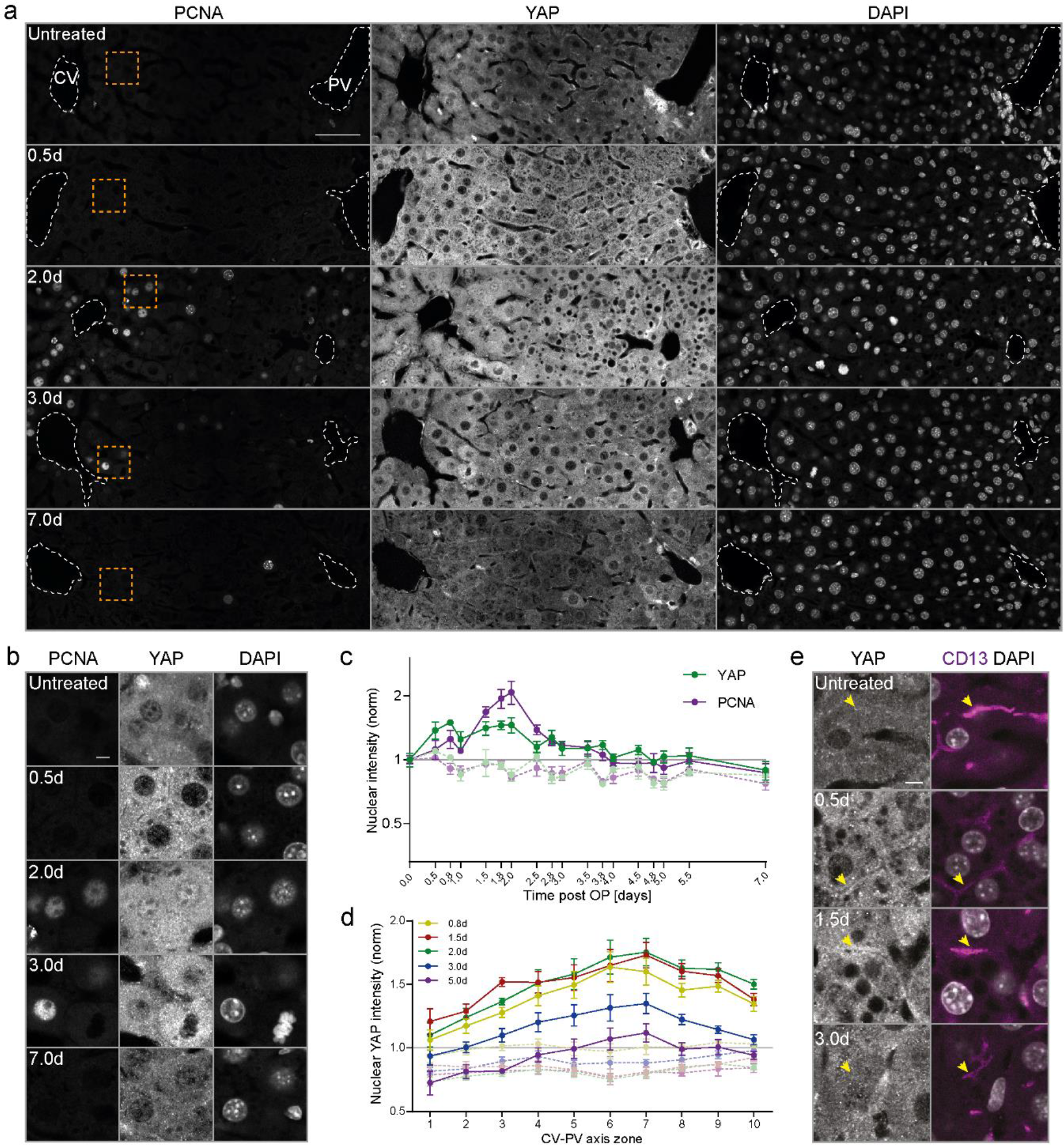
BC network expansion coincides with activation of YAP during regeneration. **a-b)** Fluorescence stainings for YAP, PCNA and with the nuclear marker DAPI on liver tissue sections from untreated mice and animals at indicated time points post PH. Images show an entire CV-PV axis (CV, left; PV, right). Indicated regions (orange squares) are shown as magnifications in (b). Note, the bright YAP fluorescence in the PV area stem from cholangiocytes of the bile duct. **c)** Quantification of the mean nuclear YAP (green) and PCNA (magenta) intensity from images of liver tissue sections at indicated timepoints post PH (solid line) and sham OP (dashed line) as representatively shown in (a) and Fig.S4a. Data was normalized to untreated animals (timepoint 0). Mean ± s.e.m, n=3-5 mice per timepoint. Nuclear YAP and PCNA intensity of PH vs. sham time course, p < 0.0001. **d)** Spatial analysis of the mean nuclear YAP intensity from images as representatively shown in (a) and Fig.S4a, at indicated time points post PH (solid line) or sham OP (dashed line) in 10 zones within the CV-PV axis (zone 1, CV area; zone 10, PV area). Data is normalized to untreated animals (not shown). Mean ± s.e.m, n=3-5 mice per timepoint. Nuclear YAP intensity of sham vs. PH mice, p < 0.0001 (at 0.8 d, 1.5 d, 2 d, 3 d) and p < 0.01 (at 5 d). **e)** Fluorescence stainings for YAP, CD13 and with the nuclear marker DAPI on liver tissue sections from an untreated mouse or animals at indicated time points post PH. Arrows indicate BC. Note the enrichment of apical YAP at 0.5 and 1.5d post PH. Images in a, b and e are background-subtracted. Scale bars, 50 µm (a, e), 5 µm (b).

### YAP localizes to apical F-actin enriched areas of hepatocytes during regeneration

To verify the apical localization of YAP, we visualized it by immunogold labelling and electron microscopy (EM) at 1.5d and 1.8d post PH (Fig.4a). Consistent with the IF, YAP was enriched at microvilli and in the sub-apical region of hepatocytes (Fig.4a’) as compared to the basolateral area (Fig.4a’’). To quantify the enrichment, we determined YAP density at the apical and basolateral membrane of hepatocytes from EM images (6×6 images, ∼30×30µm) (Fig.S5b). The density of immunogold particles was on average 4.2-fold higher in the apical than the basolateral region (Fig.S5c), demonstrating that YAP is enriched at the apical compartment during regeneration.

**Figure 4.**
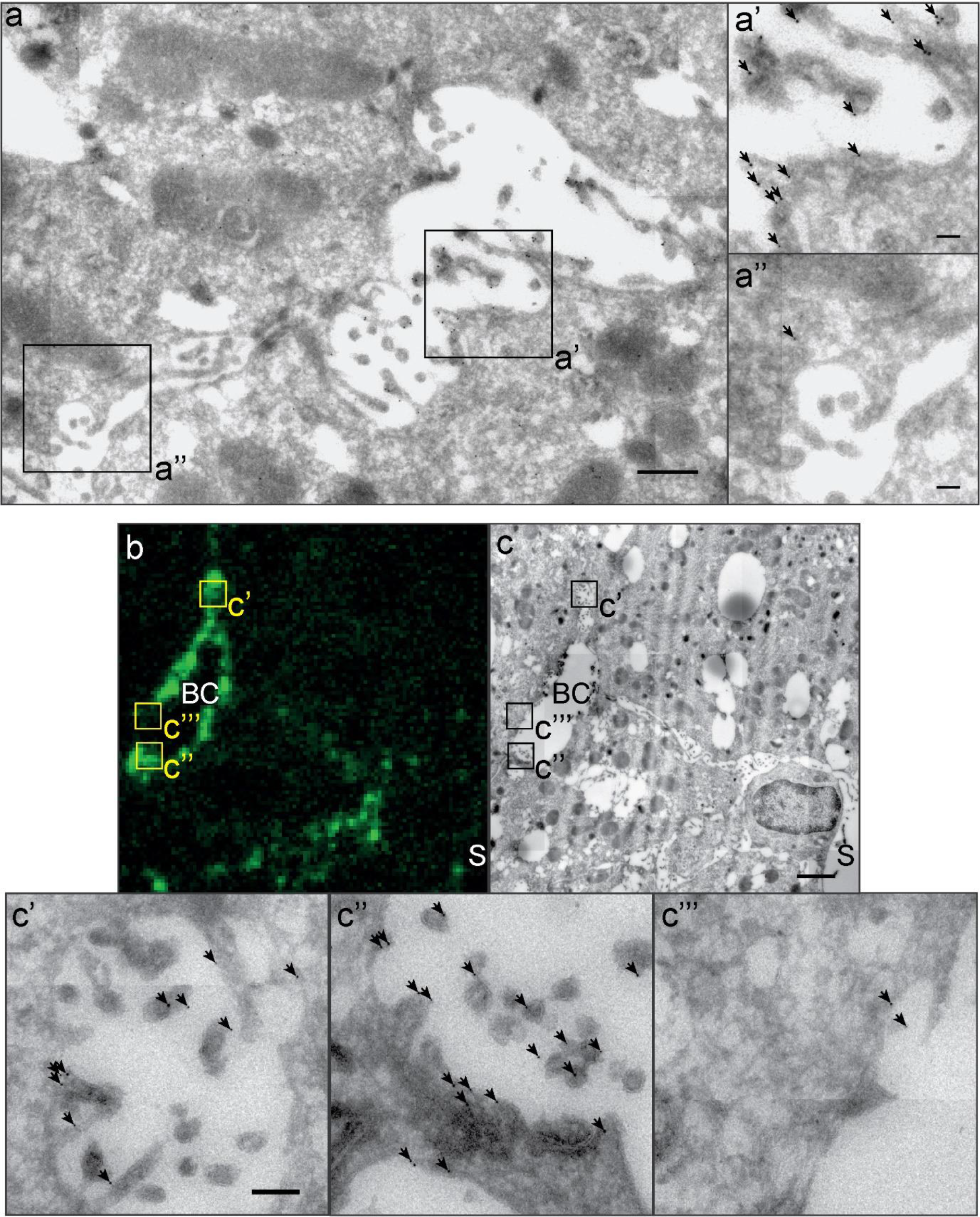
YAP enriches in F-actin rich areas in the apical compartment of hepatocytes during regeneration. **a)** EM image of immunogold-labelled YAP on liver tissue sections at 1.8 d post PH. Image (a) shows a BC in between two hepatocytes. Indicated regions (black squares) of the apical (a’) and the basolateral (a’’) membrane of the hepatocyte are shown as magnification. Arrows indicate gold particles. **b, c)** Correlative light and electron microcopy of F-actin (b) and YAP (c) on a liver tissue section at 1.8 d post PH. YAP was detected by immunogold-labelling, F-actin by fluorescence staining with phalloidin. Indicated BC regions with high (c’, c’’) or low (c’’’) F-actin levels are shown as magnifications. Arrows indicate gold particles. The image in (b) is background-subtracted. Scale bar, 0.5 µm (a), 0.1 µm (a’, a’’), 2 µm (c), 0.2 µm (c’). BC, bile canaliculus; S, sinusoid.

YAP is a mechano-sensor that responds to alterations of the actin cytoskeleton (36). From this perspective, its localization to the apical region of hepatocytes is ideal to sense alterations of biliary fluid dynamics and BC network expansion through changes of the acto-myosin system. As a pre-requisite, YAP should be enriched in areas of high F-actin content. To test for this, we used correlative light EM (CLEM) on liver tissue sections at 1.8d post PH (Fig.4b, c). F-actin was imaged at 186nm resolution to identify F-actin-rich areas, whereas YAP was imaged at 1.1-2.6nm resolution to visualize immunogold labelling. YAP was particularly enriched in the F-actin-dense sub-apical region (Fig.4c’, c’’, high F-actin levels; Fig.4c’’’, low F-actin level, see also Fig.S6). Gold particles were often associated with the microvilli in the BC, supporting the idea that YAP associates with the apical actin cytoskeleton. These results provide the first IF and EM detection of YAP at the apical compartment of hepatocytes *in vivo* and provide support to the idea that the actin-dependent mechano-sensory function of Hippo may link BA metabolism to growth control during liver regeneration.

### Bile acid-induced activation of YAP is dependent on the acto-myosin system

Given that YAP localizes to apical F-actin-rich areas in hepatocytes of regenerative liver, we hypothesized that BA may activate YAP through modulation of the acto-myosin system. We tested this idea on primary mouse hepatocytes. When cultured in collagen sandwich, these cells re-polarize forming BC *in vitro* (37) and exhibit a cholestatic-like phenotype (38), mimicking aspects of the metabolic state of the regenerative liver. Remarkably, YAP was not only cytoplasmic but also enriched at the apical region of the hepatocytes *in vitro* (Fig. 5a, upper left), as observed during regeneration *in vivo* (see Fig.3e and Fig.4). To recapitulate more closely the state of hepatocytes in the regenerating liver, we added the BA deoxycholic acid (DCA) at 200µM (Fig.5a, lower left), a concentration that is comparable to serum levels in mice after PH (39). DCA was sufficient to stimulate YAP nuclear translocation (1.7-fold increase as compared to DMSO control cells, Fig.5b), consistent with earlier observations (17). DCA also induced a strong dilation of BC (Fig.5a, compare Control vs. DCA), and up-regulation of YAP (Fig.S7a), as observed during regeneration *in vivo* (Fig.S4c). Thus, the *in vitro* system recapitulates properties of hepatocytes in the regenerating liver and, thus, proves suitable for studying YAP regulation.

**Figure 5.**
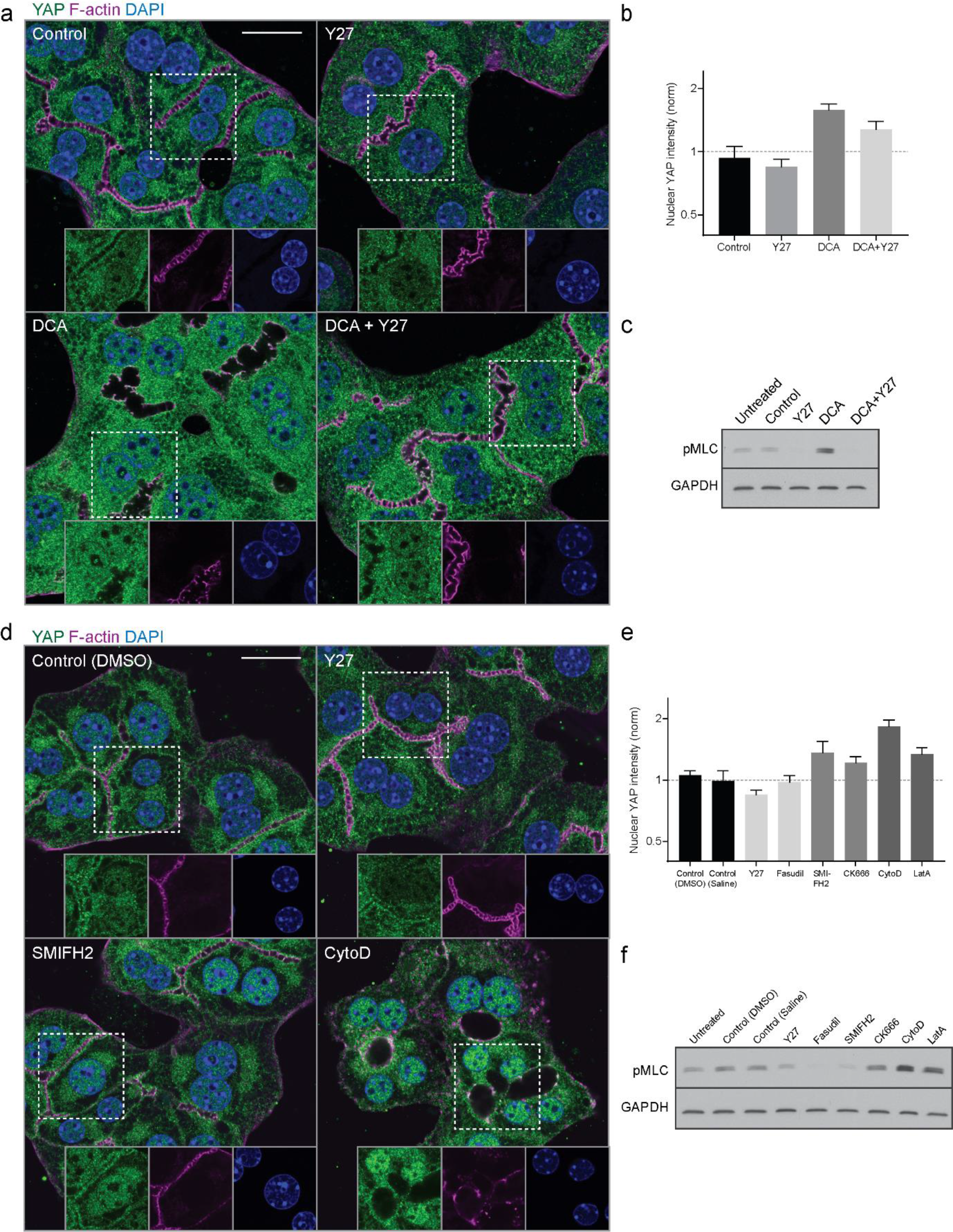
Bile acids activate YAP via the acto-myosin system. **a)** Fluorescence stainings for YAP (green), F-actin (magenta) and with the nuclear marker DAPI (blue) of primary hepatocyte cultures treated with DMSO (control), Y27, deoxycholic acid (DCA) or DCA+Y27 for ∼18 h. Indicated areas (dashed rectangle) are shown as magnifications in insets. **b)** Quantification of the mean nuclear YAP intensity from fluorescence images of primary hepatocytes treated with DMSO (control), Y27, DCA or DCA+Y27 for ∼18 h as representatively shown in (a). Data is normalized to untreated cells (not shown). Mean ± s.e.m., n = 7; DMSO vs. DCA, p < 0.0001; DMSO vs. Y27, p > 0.05 (n.s.); DMSO vs. DCA+Y27, p < 0.05; DCA vs. DCA + Y27, p = 0.05. **c**) Western blot of pMLC and GAPDH (loading control) in primary hepatocyte culture lysates. Cells were untreated or incubated for 18 h with the indicated compounds **d)** Fluorescence stainings of primary hepatocytes for YAP (green), F-actin (magenta) and with the nuclear marker DAPI (blue). Cells were treated with DMSO (control), Y27, SMIFH2 or Cytochalasin D (CytoD) for 6 h. Indicated areas (dashed rectangle) are shown as magnifications in insets. **e)** Quantification of the mean nuclear YAP intensity from images of primary hepatocytes treated with DMSO, Saline, Y27, Fasudil, SMIFH2, CK666, CytoD or Latrunculin A (LatA) for 6 h. Saline serves as control for Fasudil, DMSO serves as control for all other conditions. Inhibitors affecting similar actin processes are displayed in the same grey level. Saline, Fasudil, CK666 and LatA conditions are not shown in (d). Data is normalized to untreated cells (not shown). Mean ± s.e.m., n=3-5; DMSO vs. Y27, p < 0.01; DMSO vs. CytoD, p < 0.0001; DMSO vs. LatA, p < 0.05; all other conditions are n.s. (p > 0.05) compared to the control. **f)** Western blot of of pMLC and GAPDH (loading control) in untreated or actin inhibitor-treated primary hepatocyte culture lysates. Inhibitor treatments are the same as in (e). Scale bars, 20 µm (a, d).

We hypothesize that the stimulation of YAP nuclear translocation by DCA may be mediated by an actin-dependent mechano-sensory function. A test is whether inhibition of acto-myosin contractility abrogates the stimulatory effect by DCA on YAP nuclear translocation. Incubation of hepatocytes with the Rho kinase inhibitor Y27 blocked the activation of YAP (Fig.5a, lower right) and reduced the effect of DCA by 47% (Fig.5b). To confirm that DCA acts via the acto-myosin system, we analyzed pMLC levels in primary hepatocytes upon BA treatment by immuno-blot analysis (Fig.5c). DCA increased pMLC levels 1.57-fold as compared to untreated cells and Y27 blocked this induction (Fig.5c, Fig. S7b). This supports the hypothesis that YAP senses BA through an acto-myosin-dependent mechanism.

YAP is sensitive to perturbations of most actin cytoskeleton processes, including e.g. actin nucleation, polymerization and contractility (36,40). To verify that YAP specifically responds to alterations of acto-myosin contractility, but not to general perturbations of the actin cytoskeleton, we compared the effects of a set of small molecule inhibitors targeting distinct actin properties, (i) acto-myosin contractility using the Rho kinase inhibitors Y27 and Fasudil, (ii) actin nucleation and branching with the Arp2/3 inhibitor CK666 and the formin inhibitor SMIFH2, and (iii) F-actin polymerization using Latrunculin A and Cytochalasin D (Fig.5d). With the exception of Fasudil, which significantly reduced nuclear YAP levels as Y27, all actin inhibitors induced nuclear YAP translocation (Fig.5e), mimicking the effect of DCA.

If YAP senses contractility, as hypothesized, YAP-activating compounds should also induce contractility. Quantification of pMLC levels of hepatocytes treated with the actin inhibitors confirmed this (Fig.5f, Fig.S7c). As expected, Y27 and Fasudil reduced pMLC levels in comparison to the control. In contrast, the actin nucleation inhibitor CK666 and the polymerization inhibitors Cytochalasin D and Latrunculin A increased pMLC levels. Cytochalasin D and Latrunculin A, which caused a pronounced activation of YAP (see Fig.5e), produced the strongest increase in pMLC levels as compared to control cells (2.1-3.0-fold increase, Fig.S7c). Altogether, our *in vivo* and *in vitro* results suggest that YAP senses BA overload through changes of BC network structure via the acto-myosin-dependent regulation of the Hippo pathway.

### YAP is regulated by the acto-myosin system during regeneration

To validate our findings *in vivo*, we tested the effect of rho kinase inhibition on YAP activity during liver regeneration. We chose Fasudil, which efficiently inhibits BC contractility and bile flow *in vivo* (18) and proved to be the most potent inhibitor of MLC phosphorylation in hepatocytes *in vitro* (see Fig.S7c). Fasudil was applied by intraperitoneal injection to mice at 2d post PH or sham operation for 1 h. First, we verified the effect of Fasudil on acto-myosin activity by measuring BC diameter (Fig.6a). Spatial analysis of BC diameter showed that Fasudil strongly dilated the BC diameter in mice after PH (15.6%, zone 18) and moderately in the liver of sham OP mice (9.3%, zone 15; Fig.6b). This is consistent with the higher BC acto-myosin activity in the regenerating than control liver (Fig.2). Next, we tested the effect of Fasudil on nuclear YAP levels during regeneration (Fig.6c). Fasudil reduced nuclear YAP levels during regeneration by up to 52% as compared to control mice (Fig.6d, red lines). Note that the sham operation alone caused a small increase in nuclear YAP levels that was also reduced by Fasudil (Fig.6d, green lines). The results argue that YAP is indeed activated by acto-myosin activity during regeneration.

**Figure 6.**
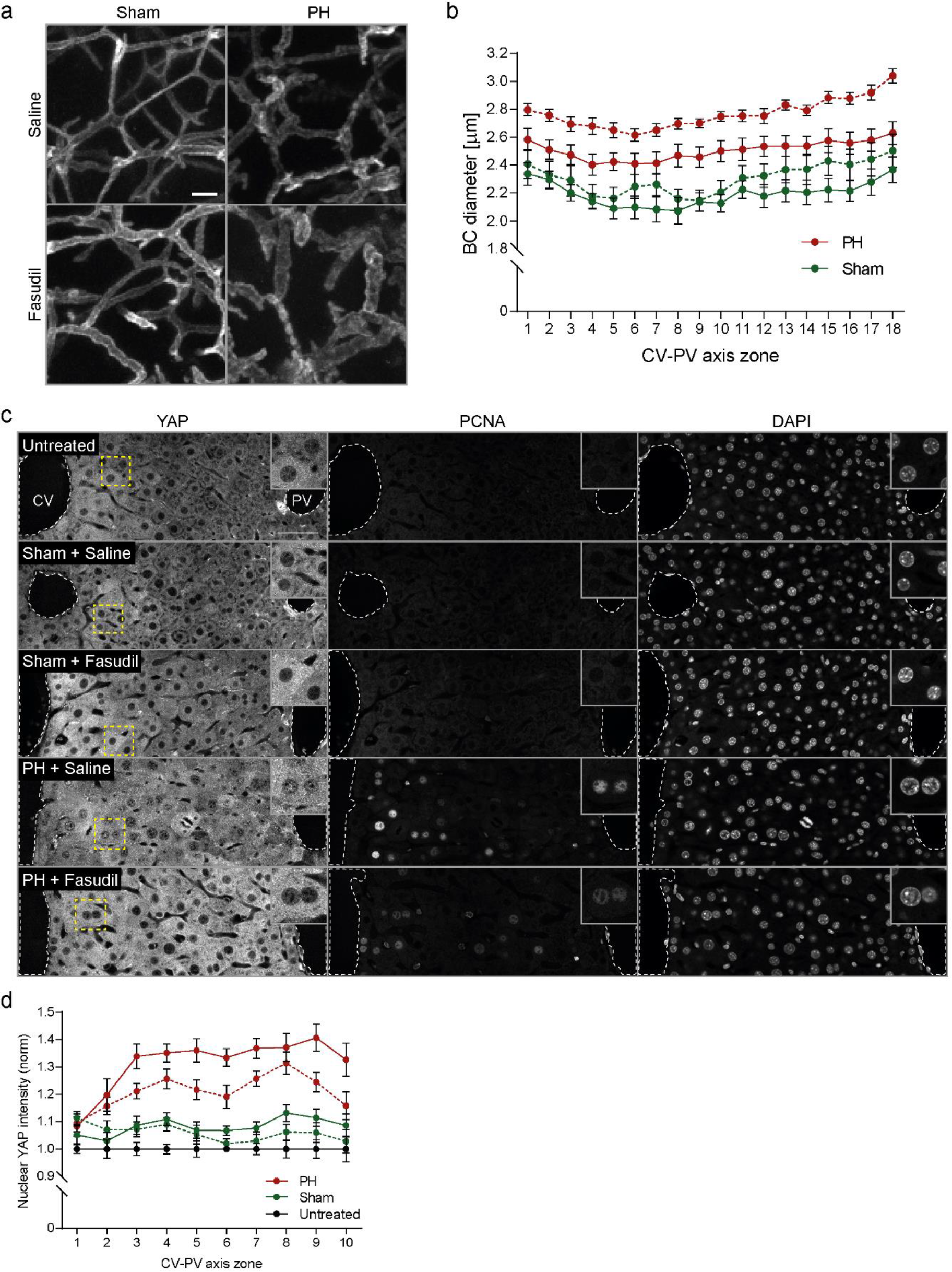
Inhibition of Rho kinase reduces YAP nuclear accumulation during regeneration. **a)** Fluorescence stainings for the apical marker CD13 on liver tissue sections from mice at 2 d post sham OP or PH, treated with saline (control) or the Rho kinase inhibitor Fasudil for 1 h. Shown are maximum projections of 50 µm stacks in the PV area. **b)** Quantification of BC diameter within 18 zones along the CV-PV axis (zone 1, peri-central; zone 18, peri-portal) from mice at 2 d post sham OP (green) or PH (red), treated with saline (control, solid line) or Fasudil (dashed line). Diameter was measured from 3D BC network reconstructions of image stacks as representatively shown in (a). The zones directly adjacent to the CV and PV were excluded from the analysis (∼ 1 cell layer). Mean ± s.e.m, n=7-8 mice per condition from 3 independent experiments. BC diameter of sham operated mice treated with saline vs. Fasudil, p < 0.0001; BC diameter of PH mice treated with saline vs. Fasudil, p < 0.0001. **c)** Fluorescence stainings for YAP, PCNA and with the nuclear marker DAPI on liver tissue sections from untreated mice or animals at 2 d post sham OP or PH, treated with saline (control) or Fasudil for 1 h. Indicated regions (dashed rectangle) are shown as magnifications in insets. **d)** Quantification of the mean nuclear YAP intensity within 10 zones along the CV-PV axis, (zone 1, peri-central; zone 10, peri-portal) from IF images as representatively shown in (c). Data was normalized to untreated animals. Mean ± s.e.m, n=7-8 mice per condition from 3 independent experiments. Nuclear YAP intensity of sham operated mice treated with saline vs. Fasudil, p > 0.05 (n.s.); nuclear YAP intensity of PH mice treated with saline vs. Fasudil, p < 0.001. Scale bars, 5 µm (a), 50 µm (c).

### A mechanistic model predicts a switch-like activation of YAP upon mechanical stimulation during regeneration

Large and sustained increases in BA loads, as after PH, stimulate liver re-growth (9). In contrast, moderate fluctuations of BA concentrations, resulting e.g. from circadian rhythm or diet (41,42), can be buffered by the metabolic activity of hepatocytes and do not result in YAP activation and a proliferative response. This raises the question of how YAP can discriminate between physiological changes in BA loads and pressure in the BC versus those that require cell proliferation.

Bile flow through bile canaliculi is driven by both the osmotic pressure of actively pumped bile salts and osmolytes, and contractility of BC (18,27). Upon PH without removal of the gall bladder, the total bile salt pool in the body is only marginally reduced, since intra-hepatic BA account for 2-4% of the total pool (43). Therefore, the liver remnant needs to transport the full bile salt pool through a reduced BC network, leading to an increased bile salt flux per liver weight (44) and, consequently, increased osmotic pressure and acto-myosin tension in the BC. It is currently technically impossible to measure pressure within the BC. However, it is possible to predict the changes in apical cortical tension and couple these to YAP nuclear translocation using a theoretical approach. Based on our experimentally measured spatial profile of BC diameter (see Fig.1b), we developed a biophysical model (see Supplemental Experimental Procedures) that predicts the apical cortical tension during regeneration from the osmotic pressure within the BC network. The model predicts a ∼2-fold increase of apical tension throughout the entire CV-PV axis (Fig.S8a) as an immediate response to liver resection within 0.8d. Comparison of the inferred cortical tension and our experimentally measured apical pMLC levels shows a high correlation (Pearson correlation r=0.94, Fig.S8b), supporting the predictive power of the model.

Next, we coupled the biophysical model with a biochemical model of YAP regulation to predict nuclear YAP levels from cortical tension. Based on reports on the regulation of YAP in the liver (11,13,45), we considered five regulatory mechanisms: YAP synthesis, degradation, phosphorylation, cytoplasmic sequestration as well as nuclear-cytoplasmic shuttling (Fig.7a). Assuming simple mass action and Michaelis-Menten kinetics for the considered reactions (see Fig.7a), the model has 16 parameters. Of these, the compartment volumes of hepatocyte nuclei and cytoplasm were experimentally measured (19) (see Table S2). Five parameters were set to the value 1, as the analytical analysis of the model revealed that they do not affect the steady state solution. The remaining 9 parameters were estimated by fitting the quasi steady state solution of the model to 120 data points (nuclear and total YAP measurements; for details see Supplemental Experimental Procedures and Table S2). To test the model, we predicted the nuclear and total YAP levels from the estimated apical cortical tension and compared the results to our experimental measurements (Fig.S8c, d). Despite its relative simplicity, our mechanistic model reproduced the experimentally determined values of nuclear and total YAP levels over the time course of regeneration.

**Figure 7.**
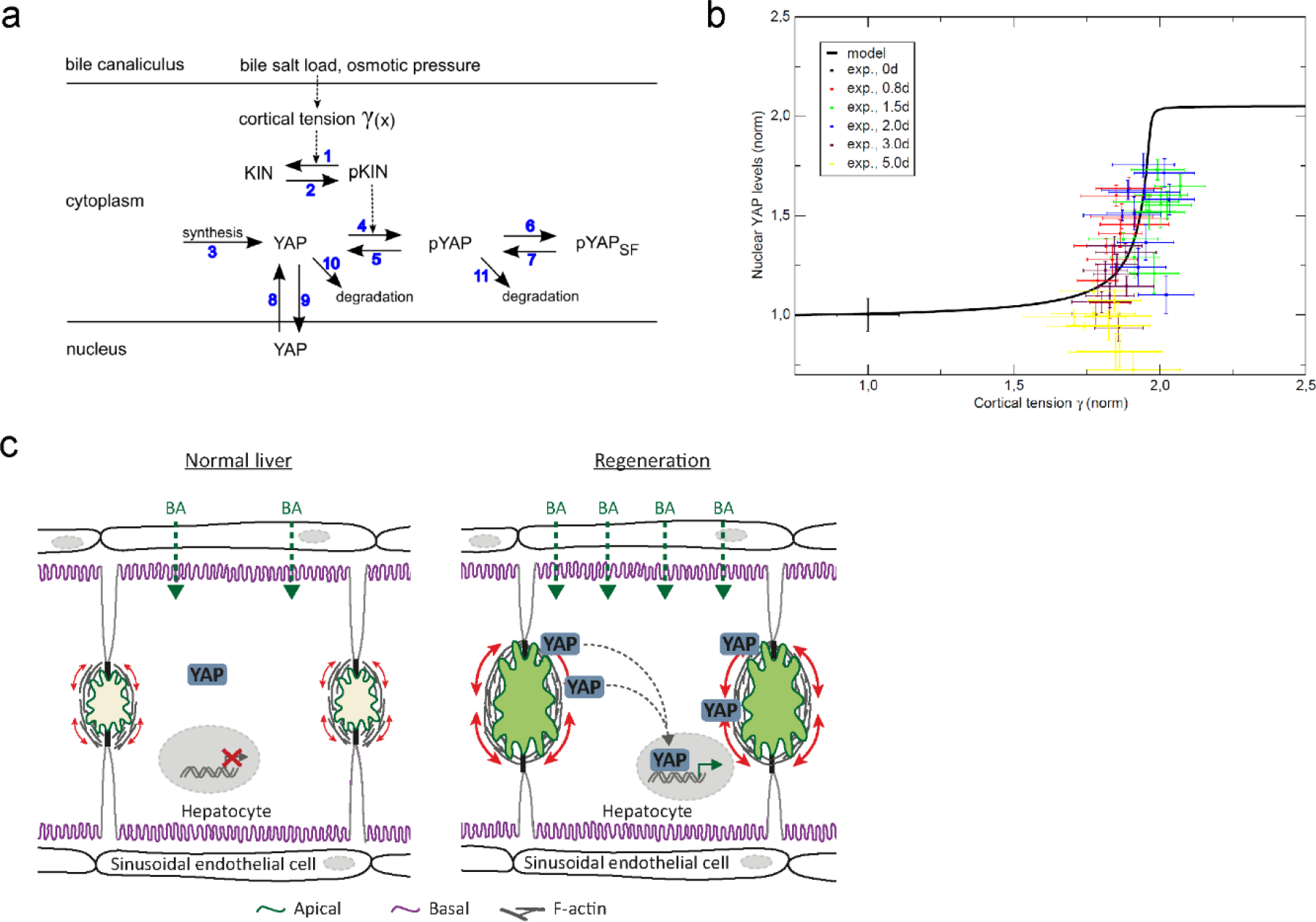
Model of mechano-sensing of metabolic status during liver regeneration. **a**) Model of YAP regulation by cortical tension of the BC. Arrows denote reactions or transport steps. Blue numbers label individual processes and associated parameters as listed in Table S2. KIN, kinase; pKIN, phosphorylated kinase; pYAP, phosphorylated YAP; SF, sequestration factor. See Experimental Procedures for model equations and analysis. **b**) Model prediction (solid black line) of cortical tension and nuclear YAP levels reveals a sigmoidal stimulus-response curve. Experimental data (symbols) of nuclear YAP is reproduced from Fig.3d and mapped to cortical tension levels using experimental data from Fig.1b and Fig.2d. **c**) Schematic drawing of YAP regulation by BA through canalicular cortical tension. Tissue resection by partial hepatectomy induces a BA overload which elevates osmotic pressure and cortical tension of bile canaliculi. Concomitant changes of apical acto-myosin properties recruit YAP to the apical cortex where it is activated for nuclear translocation.

Finally, we used the model to infer the stimulus-response behavior, by analyzing YAP translocation to the nucleus as a function of BC cortical tension. The resulting stimulus-response curve (Fig.7b, black curve) is robust to changes of the kinetic rate constants. The nuclear YAP measured values match the theoretical curve very well (Fig.7b, colored symbols). The stimulus-response curve reveals a sharp threshold-like sigmoidal dependency where YAP translocates into the nucleus only if the relative apical cortical tension exceeds ∼1.75-fold (Fig.7b). This suggests that low and moderate fluctuations of BA load and cortical tension (e.g. diet-triggered) are tolerated and do not trigger the Hippo pathway. In contrast, severe alterations of biliary pressure, such as after PH, robustly activate YAP in a switch-like manner to induce regeneration. Our model therefore predicts the existence of a threshold in the activation of YAP to ensure a switch from a non-proliferative to a proliferative response.

## Discussion

We described a novel mechanism whereby hepatocytes detect organ size during liver regeneration based on mechano-sensing of metabolic load through the apical acto-myosin system (Fig.7c). We demonstrate that liver resection induces structural and functional changes of the BC network, including expansion of the apical surface of hepatocytes and increase in acto-myosin contractility. These changes occur as a compensatory response to BA overload induced by tissue resection, and are sensed by YAP, which is enriched at the apical domain of hepatocytes and translocates to the nucleus dependent on the activity of the apical acto-myosin system. During regeneration, the restoration of the liver-to-body weight ratio also re-establishes the metabolic homeostasis, the structure of the BC network reverts to normal, and YAP returns to its physiological state (cytoplasmic localization). Thus, the BC network represents a self-regulatory mechano-sensory system that adapts to the overall metabolic demand of the body and acts as readout of tissue status.

The function of BA as regulators of liver regeneration has been demonstrated early on, and mainly attributed to their signaling function via nuclear receptors (9). BA metabolism is a common target in clinical practice for treatment of a variety of human diseases to promote liver function. Whereas elevated BA levels promote hepatocyte proliferation, deprivation of BA results in a regenerative delay (9). However, besides signal messengers, BA are also regulators of the Hippo pathway, suggesting the existence of an additional mechanism (17). Our results implicate the BC network in such a mechanism, by playing a mechano-sensory role, linking BA metabolism, liver tissue structure and Hippo signaling. A remarkable feature of liver tissue organization is that all hepatocytes are connected via the BC, bridging the sub-cellular to the tissue level. Such organization permits the communication of global tissue properties, e.g. metabolic status, to each individual cell within the lobule, to control cell behavior collectively. Key for the signaling function of the BC network is the dynamic nature of the apical plasma membrane of hepatocytes, which can promptly respond to metabolic changes. Bile pressure forms a gradient within the lobule varying ∼30-fold from the CV to the PV area (18). Such variation needs to be balanced by the tension of the acto-myosin system. This implies that changes of pressure need to be sensed, evoking the need for a mechanosensory system. The same system may conceivably operate during liver regeneration.

Our mathematical model predicts a sharp pressure-dependent threshold in the activation of YAP to ensure a robust switch from a non-proliferative to a proliferative response to the metabolic overload. This means that fluctuations of BA concentrations in the physiological range could be compensated by the metabolic activity of hepatocytes, whereas large and sustained changes, as after PH, trigger the Hippo pathway in a switch-like manner to induce regeneration. Our mechanistic mathematical model of signal transduction through the Hippo pathway triggered by a mechanical stimulus may be applicable to organ size control more generally, e.g. growth of the fly wing imaginal disc in response to apical membrane tension (46).

A wide array of apical actin interacting and polarity or cell-junction proteins have already been reported to regulate the Hippo pathway (16,47). All these molecules are potential YAP regulators during liver regeneration. Given the collective properties of the apical plasma membrane and associated acto-myosin cytoskeleton, a systems analysis of its complex super-molecular network will be required to identify the precise underlying molecular mechanisms.

## Experimental Procedures

### Animal work

Animal experiments were performed on 8-12 weeks old, male, C57BL/6JOlaHsd (Harlan laboratories) mice at the MPI-CBG (Dresden, Germany). PH and BDL experiments were performed between ∼8am–1pm. Experiments were conducted in accordance with German animal welfare legislation and in strict pathogen-free conditions in the animal facility of the MPI-CBG. Protocols were approved by the Institutional Animal Welfare Officer (Tierschutzbeauftragter) and all necessary licenses were obtained from the regional Ethical Commission for Animal Experimentation of Dresden, Germany (Tierversuchskommission, Landesdirektion Dresden). Mice were fasted 6h prior sacrifice (water ad libitum).

### PH

PH was performed based on (20) but without removal of the gall bladder (left and right median lobes resected individually). Sham operated mice received the same treatment but without liver resection.

### YAP knockout

A conditional-hepatocyte specific mosaic YAP knockout was induced by adeno-associated virus mediated expression of iCre recombinase using AAV/DJ-pALB(1.9)-iCre (Vector biolabs, USA) in YAP^fl/fl^ mice (21). AAV/DJ-pALB(1.9)-eGFP (Vector biolabs, USA) served as control.

### Fasudil administration

Mice received 30mg/kg Fasudil or saline intravenously and were sacrificed 1h after.

### Bile duct ligation

BDL was performed as described in (22) and sacrificed 1d post-surgery. Sham operated mice received the same treatment without BDL.

### Tissue fixation

Anaesthetized mice were perfused transcardially with 4% PFA/0.1% Tween20/PBS. Tissue was post-fixed in 4% PFA/0.1% Tween20/PBS for ∼48h at 4°C for IF or in 1% glutaraldehyde/3% PFA/PBS for 1.5h for morphological EM.

### Primary hepatocytes

Hepatocytes were isolated and cultures as described in (23) and (24), respectively. Cells were treated with 20µM Y27, Fasudil, SMIFH2, CK666 or 5µM Latrunculin A or Cytochalasin D for 6h. For BA experiments, cells were pre-incubated with 20µM Y27 or DMSO (control) for1 h, followed by 16-18h incubation with DMSO, 200µM DCA, 20µM Y27 or 20µM + 200µM DCA. Cells were lysed for biochemical analysis or fixed with 4% PFA for 30min.

### IF stainings

100µm liver slices were permeabilized with 0.5% TritonX100 for 1h, quenched with 10 mM NH4Cl for 30min, blocked with 0.2% gelatin/300mM NaCl/0.3% TX100/PBS and incubated with primary (Table S1) and fluorescently-conjugated secondary antibody in blocking buffer for 48h each. Co-staining of CD13 with PCNA and YAP or pMLC required additional antigen retrieval in citric acid or EDTA buffer at 80°C for 1h, respectively. To preserve CD13 staining during retrieval, tissue was pre-fixed with 4% PFA for 20min. Following retrieval, sections were stained with primary antibodies (Table S1) as described above. Sections were mounted in 90% glycerol (2D microscopy) or cleared with SeeDB (3D imaging) as previously described (25). IF staining of hepatocytes was performed as described previously (24).

### Microscopy

Samples were imaged on a Zeiss LSM780 confocal microscope as 2×1 image tiles covering an entire CV-PV axis as single plane image or 30 μm z-stacks (voxel size of 0.28 × 0.28 × 0.3μm) as described in (18,19).

### Western blot

SDS-PAGE separated liver lysates were transferred onto nitrocellulose membrane. Proteins were immune-detected (Table S1) using chemiluminescence. The relative density of bands was quantified using the Fiji software.

### EM

For morphological EM, 70 nm liver sections were imaged as 10×10 grids (image size 19×19µm, pixel size 9nm) on a Philips Tecnai12 EM (FEI, USA). For immuno-EM, sections were immunolabelled using YAP antibody (Table S1) and goat-anti-rabbit-10nm gold coupled IgG as previously described (26) and imaged at 1.1-2.6nm resolution. For CLEM, sections were additionally incubated with Alexa Fluor488-conjugated phalloidin and imaged at 0.186 µm/pixel resolution on a Zeiss Axioplan2 light-microscope.

### Image analysis and mathematical modeling

See supplemental experimental procedures.

## Acknowledgements

We thank the Biomedical Services (Jussi Helppi, Anne-Muench Wuttke and Barbara Langen), Light Microscopy Facility (Jan Peychl), Electron Microscopy Facility (Jean-Marc Verbavatz) of the MPI-CBG and the Centre for Information Services and High Performance Computing (ZIH) of the TU Dresden for the generous provision of computing power.

## Author Contributions

K.M. and M.Z. conceived the project and K.M. designed most of the experimental strategy. K.M., with help of S.S., conducted the experiments. M.W.-B. performed immuno-EM and CLEM. U.D. trained K.M. in PH. H.M.-N and Y.K. developed the image analysis algorithms. E.M.T provided the YAP antibody, a key reagent for the study. L.B. and Y.K. developed and analyzed the mathematical model. K.M. and M.Z. wrote the manuscript.

