## Supplemental file for "Metabolic control of YAP via the acto-myosin system during liver regeneration"

**Supplemental Information**

**Supplemental Figures**

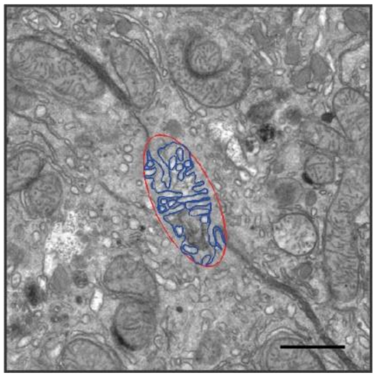

**Figure S1 Determination of BC membrane length and perimeter from EM images**
Representative EM image segmentation of the BC membrane (blue) and the minimal enclosing ellipse (perimeter) of that segmentation (red). Scale bar, 1 µm.

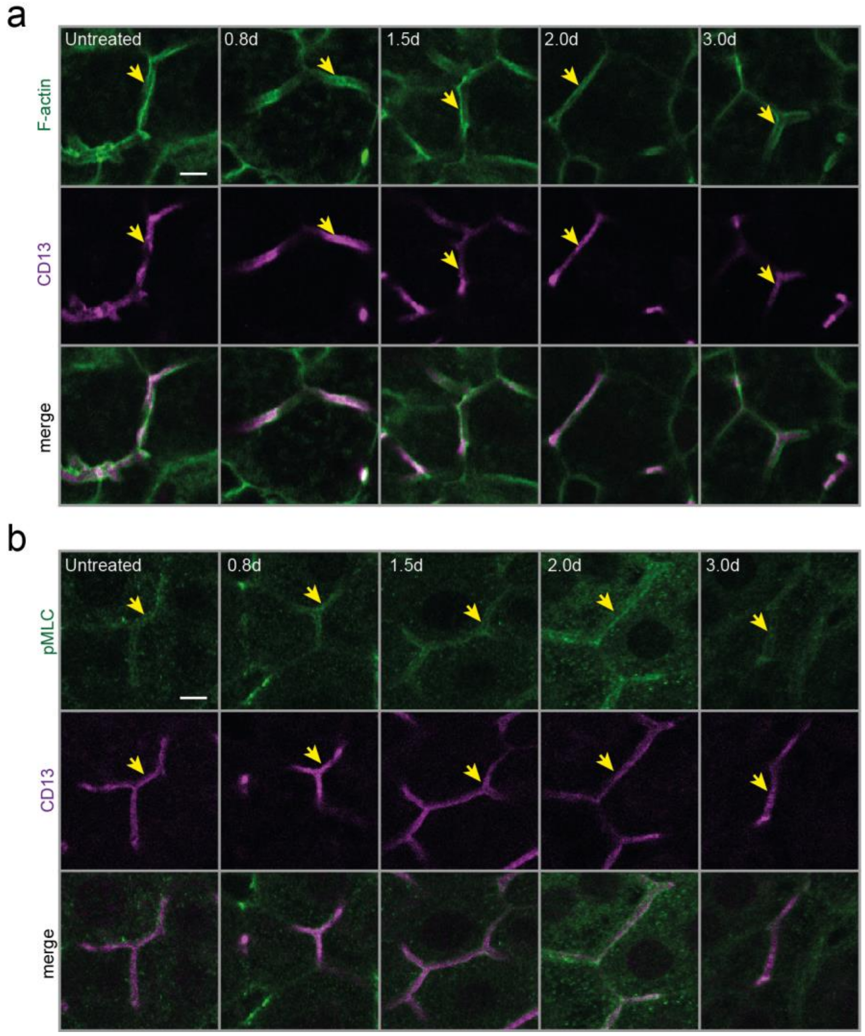


**Figure S2 Fluorescence staining of apical F-actin and pMLC in livers after sham OP**

**a, b)** Fluorescence stainings for F-actin (a) or pMLC (b) and the apical marker CD13 on liver tissue sections from untreated mice or animals at indicated time points post sham OP. Images were taken in the PV area. Arrows indicate BC. Scale bar, 5 µm (a, b).

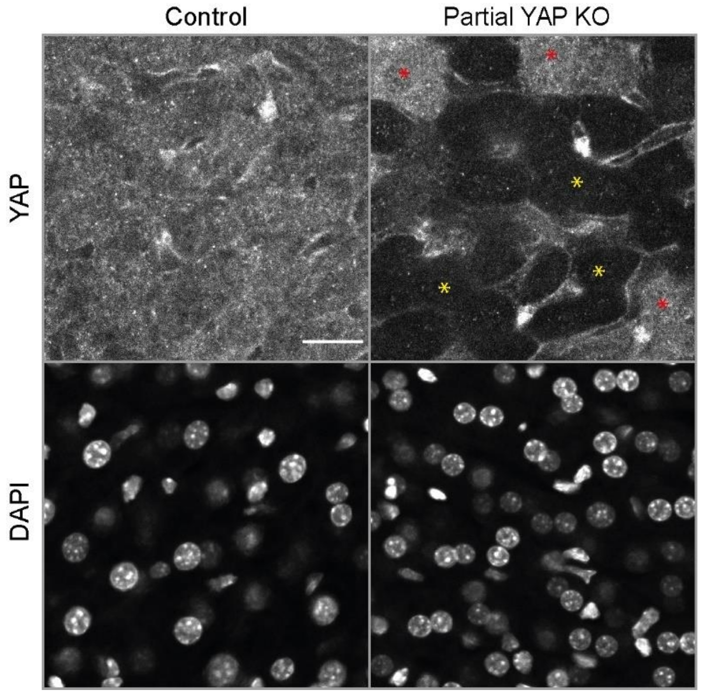


**Figure S3 Validation of YAP antibody on liver tissue sections**

Fluorescence staining for YAP and DAPI on liver tissue sections from control or conditional partial YAP knockout (KO) in mice. The knockout was specifically induced in hepatocytes by adeno-associated virus mediated expression of Cre-recombinase from an Albumin promotor (pALB) in $YAP^{fl/fl}$ mice. Yellow asterisks indicate KO hepatocytes, red asterisks indicate uninfected cells. Control mice received EGFP expressing adeno-associated-virus. Scale bar, 20 µm.

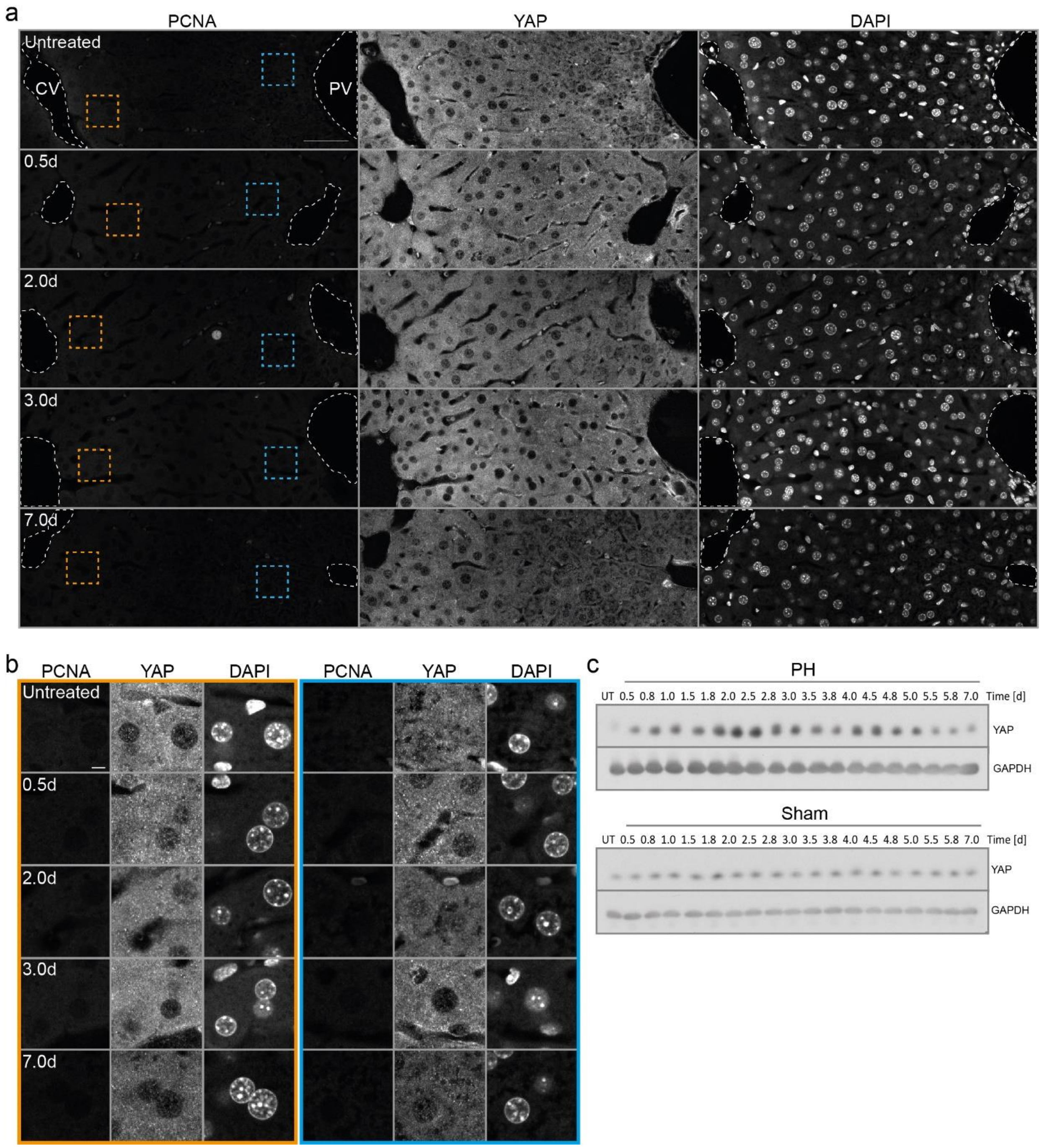


**Figure S4 Immunofluorescence staining for YAP and PCNA on liver tissue sections after sham OP**

**a, b)** Fluorescence stainings for YAP, PCNA and with the nuclear marker DAPI on liver tissue sections from untreated mice or animals at indicated time points post sham OP. Images show an entire CV-PV axis (CV, left; PV, right), veins are indicated by white dashed lines. Indicated regions (dashed rectangle) in the CV (orange) and PV (blue) area in (a) are shown as magnifications in (b). **c)** Western blot detection of YAP and GAPDH (loading control) in liver tissue lysates of untreated mice or animals at indicated time points post PH (upper panel) or sham OP (lower panel). UT, untreated control. Images in (a) and (b) are background-subtracted. Scale bars, 50 µm (a), 5 µm (b).

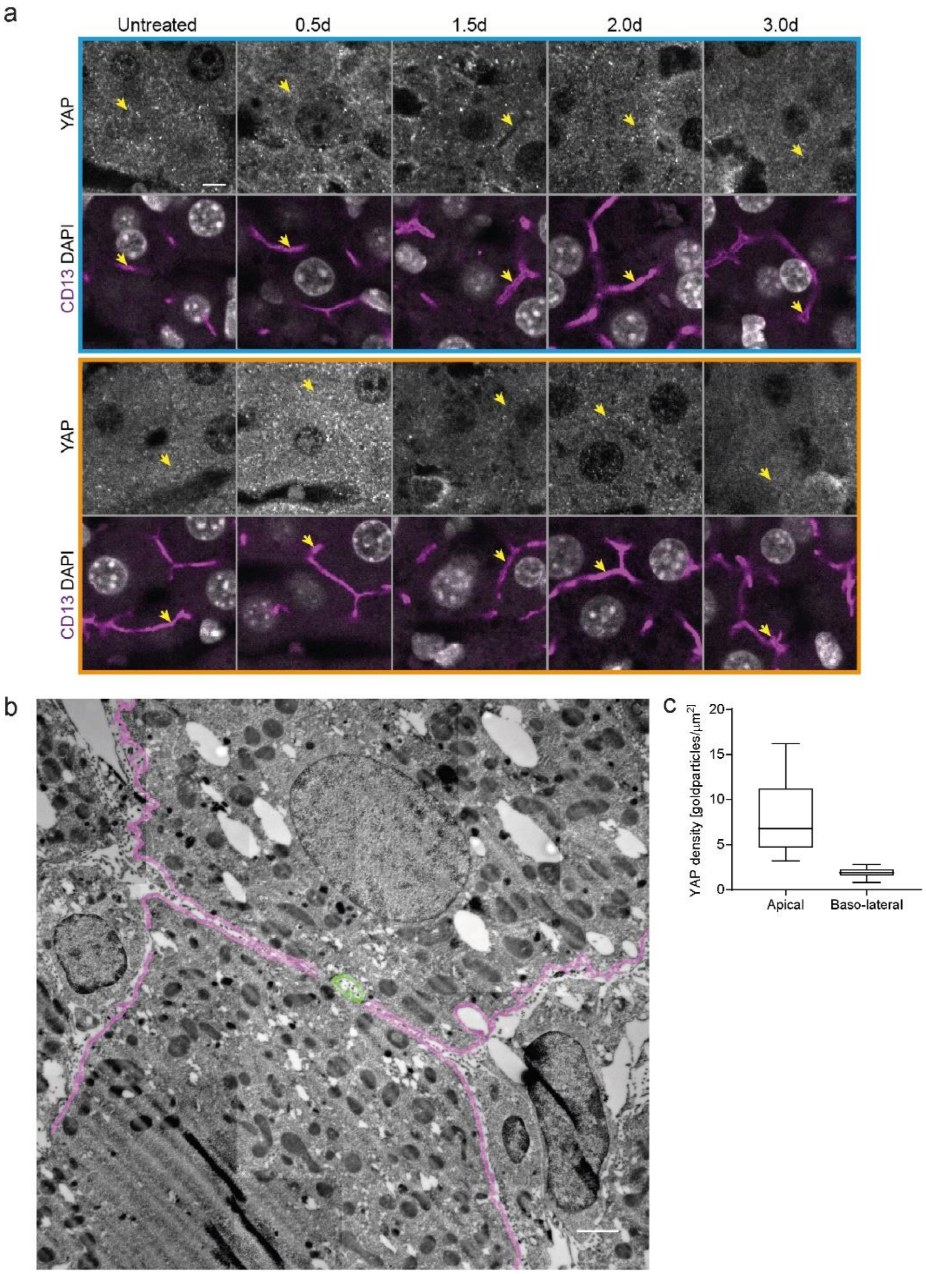


**Figure S5 Analysis of apical YAP localization**

**a)** Fluorescence stainings for YAP, CD13 and with the nuclear marker DAPI on liver tissue sections from untreated mice or animals at indicated time points post sham OP in the PV (blue, upper panel) and CV (orange, lower panel) area. Arrows indicate BC. **b)** Representative EM image showing segmentations of the sub-apical (green) and basolateral (magenta) area within a distance of 200 nm below the plasma membrane. **c)** Quantification of YAP density in the sub-apical or -basolateral membrane area (200 nm below the membrane) from YAP immuno-EM image grids on liver tissue sections at 1.5-1.8 d post PH as shown in (b). Box-whisker plot with median, 25-75 quartiles and minimum/maximum error bars, n=13 EM image grids from a total of 2 mice. Gold particle density in apical vs. basolateral area, $p < 0.0001$. Images in (a) are background-subtracted. Scale bars, 5 µm (a), 2 µm (b).

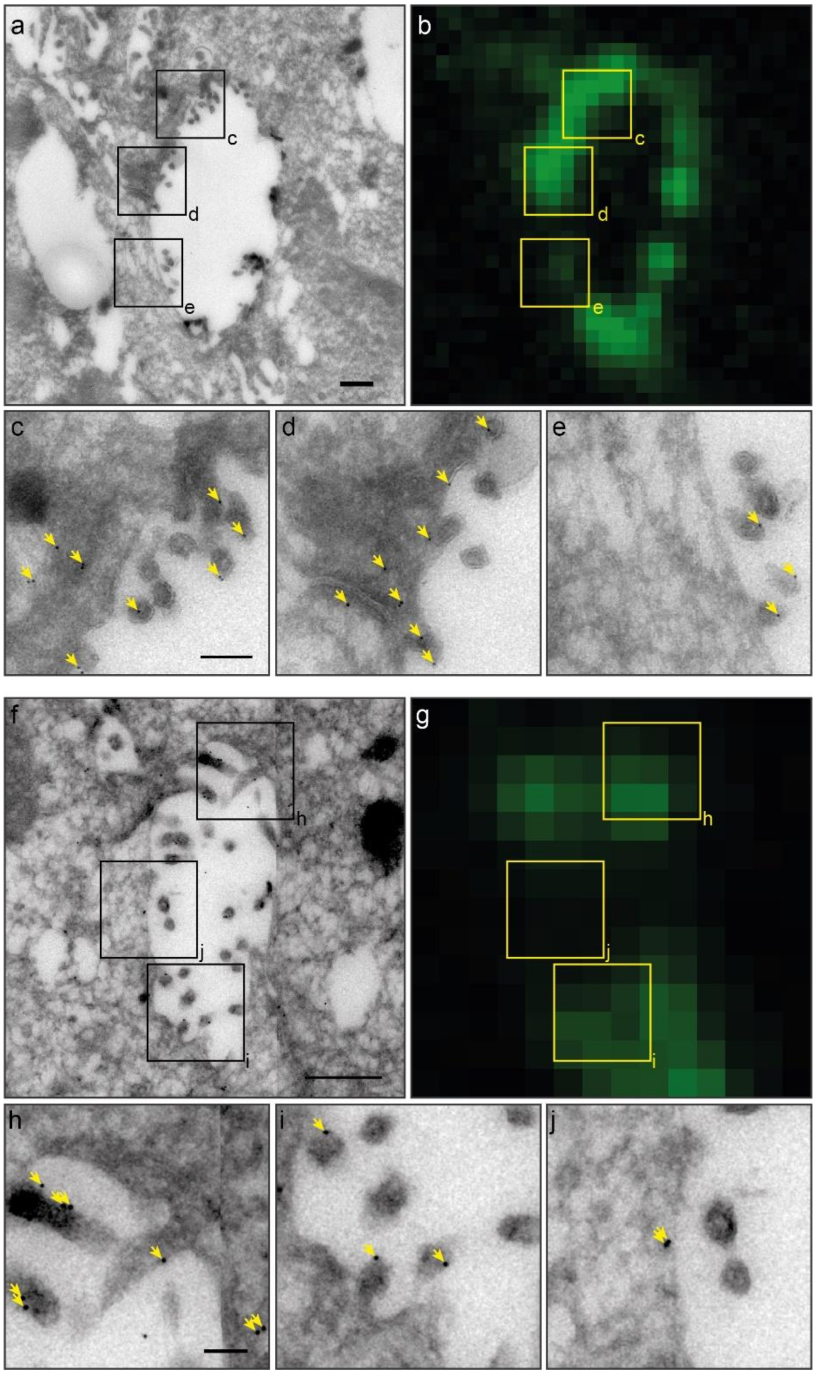


**Figure S6 Correlative light and electron microscopy of F-actin and YAP on liver tissue sections during regeneration**

**a-e)** Correlative light and electron microcopy of F-actin and YAP on a liver tissue section at 1.8 d post PH. YAP was detected by immunogold-labelling, F-actin by fluorescence staining with phalloidin. Shown are 2 BC (a, f), their respective F-actin staining (b, g) as well as magnifications of the indicated areas (black rectangle) with high (c, d and h, i) or low (e and j) F-actin levels. Arrows indicate gold particles. Images in (b) and (g) are background-subtracted. Scale bar, 0.5 µm (a, f), 0.2 µm (c), 0.1 µm (h).

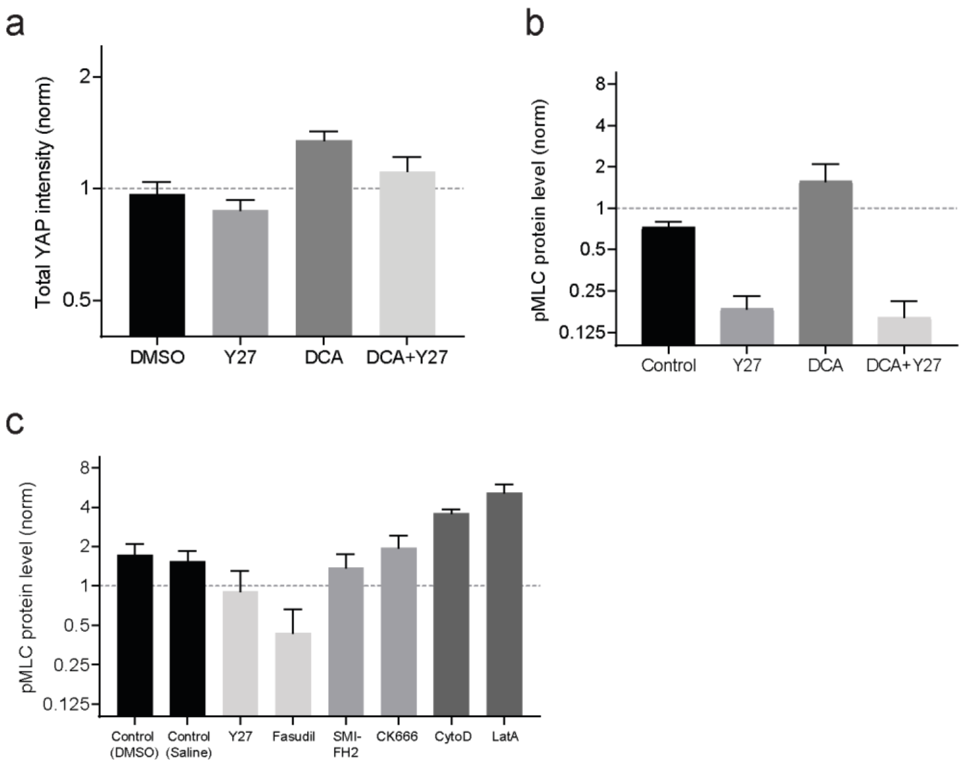


**Figure S7 Quantification of YAP and pMLC levels in primary hepatocyte cultures upon treatment with DCA and actin inhibitors**

**a**) Quantification of the mean cellular YAP levels in control (DMSO) and Y27, DCA and DCA+Y27 treated hepatocyte cultures. For full description of conditions, see legend of Fig. 5b. Data was normalized to untreated condition (not shown). Shown are mean ± s.e.m, n= 7. Total YAP intensity of DMSO vs. DCA treated cells, $p < 0.001$; DMSO vs. DCA+Y27, $p > 0.05$; DMSO vs. Y27, $p > 0.05$; DCA vs. DCA+Y27, $p > 0.05$. **b**) Quantification of pMLC in primary hepatocyte culture lysates from Western blot as representatively shown in Fig.5c. Cells were untreated or incubated for 18 h with the indicated compounds. Data is normalized to untreated cells (not shown). Mean ± s.e.m., n = 6. pMLC protein levels of DMSO vs. Y27 or DCA+Y27 treated cells, $p < 0.0001$; DMSO vs. DCA, $p = 0.11$ (n.s.). **c**) Quantification of pMLC in actin-inhibitor primary hepatocyte culture lysates from Western blot as representatively shown in Fig.5f. Cells were treated with DMSO, Saline, Y27, Fasudil, SMIFH2, CK666, CytoD or Latrunculin A (LatA) for 6 h. Saline serves as control for Fasudil, DMSO serves as control for all other conditions. Data is normalized to untreated cells (not shown). Mean ± s.e.m., n = 5. pMLC protein level of DMSO vs. CytoD treated cells, $p < 0.0001$; DMSO vs. LatA, $p < 0.001$; Saline vs. Fasudil, $p < 0.01$; all other conditions are n.s. ($p > 0.05$) compared to the control.

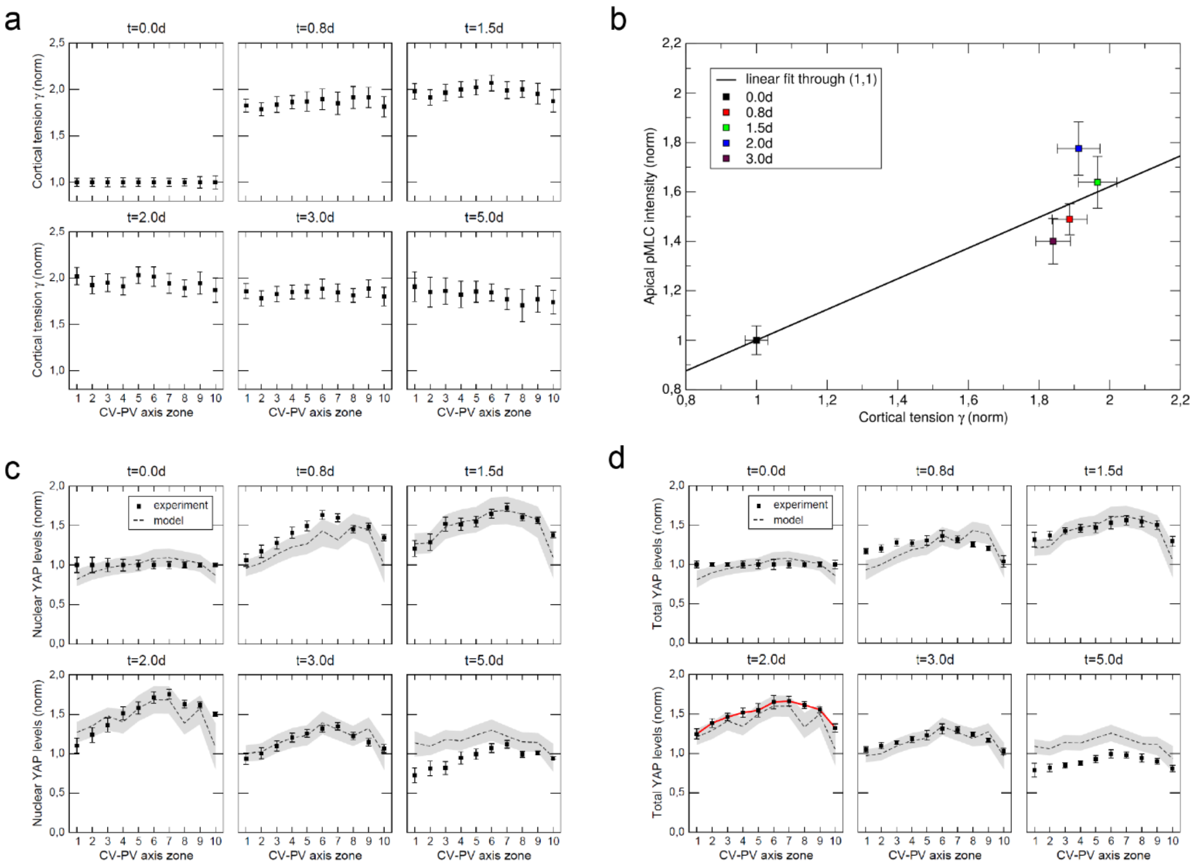


**Figure S8 Prediction of cortical tension and YAP behavior during regeneration**

**a**) Spatial profiles of predicted cortical tension during regeneration. Shown are predictions of the normalized mean ± s.e.m. cortical tension within 10 zones between the CV (zone 1) and PV (zone 10) axis at indicated timepoints (0.8 – 5.0 d post PH). The untreated condition is denoted as timepoint 0.0 d. The input data from Fig. 1b at 18 spatial positions was interpolated to align with the 10 positions of data from Fig. 3d. **b**) Correlation between predicted cortical tension and measured apical pMLC levels at different time points post PH (see legend) and in the untreated liver (t = 0.0 d). The apical pMLC intensity is derived from data reported in Fig. 2d and cortical tension from panel (a) is averaged over the corresponding positions in the PV area. The diagonal line represents the linear curve fit, see text. **c, d**) Comparison between the predicted (dashed black line) and measured (symbols) norm. nuclear (c) or total (d) YAP levels within 10 zones between the CV (zone 1) and PV (zone 10) axis at indicated timepoints (0.8 – 5.0 d) post PH and in the untreated liver (0.0 d). Shown are mean ± s.e.m. (error bar or grey corridor). In (d), the model input *s(x)* is shown as red curve (connecting the means of measured data). Experimental data in (c) is reproduced from Fig. 3d.

### Supplemental Table

*Table S1 Primary antibodies*

| Antibody | Dilution | Source |
|---|---|---|
| CD13 | 1:400 (IF) | Acris Antibodies, Germany, Cat# SM2298P |
| PCNA | 1:200 (IF) | Cell Signaling Technologies, USA, Cat# 8580 |
| YAP | 1:2000 (IF), 1:50 (EM) | Laboratory of Prof. Elly Tanaka, IMP, Vienna, Austria |
| pMLC | 1:100 (IF) | Abcam, UK, Cat# ab2480 |
| YAP | 1:1000 (WB) | Cell Signaling Technology, USA, Cat# 4912 |
| GAPDH | 1:2000 (WB) | Sigma-Aldrich, USA, Cat# G8795 |
| alpha-tubulin | 1:1000 (WB) | Sigma Aldrich, USA, Cat# T6199 |

*IF, immunofluorescence; WB, Western Blot; EM, electron microscopy*

*Table S2 Parameter values for the biochemical model of YAP regulation*

| Symbol | Name | Value | Unit | Source |
|---|---|---|---|---|
| $V_c$ | Volume of cytoplasm per cell | 5450 | $\mu m^3$ | Measured on dataset reported previously (1) |
| $V_n$ | Volume of all nuclei per cell | 860 | $\mu m^3$ | Measured on dataset reported previously (1) |
| $k_1$ | Max. flux of kinase dephos. | 1 | $1/\mu m^3$*d | free choice |
| $K_{M1}$ | M-M const. of kinase dephos. | 0.0008 | $1/\mu m^3$ | fit |
| $k_2$ | Max. flux of kinase phosphor. | 2.02 | $1/\mu m^3$*d | fit |
| $K_{M2}$ | M-M const. of kinase phos. | 0.25 | $1/\mu m^3$ | fit |
| $k_{3,0}$ | Factor of YAP synthesis rate | 1.7 | 1/d | fit |
| $k_4$ | YAP phosphorylation rate | 0.19 | 1/d | fit |
| $k_5$ | YAP dephosphorylation rate | 1 | 1/d | free choice |
| $k_6$ | pYAP binding rate to SF | 0.18 | 1/d | fit |
| $k_7$ | pYAP unbin. rate from SF | 1 | 1/d | free choice |
| $k_8$ | YAP export rate from nucleus | 1 | 1/d | free choice |
| $k_9$ | YAP import rate into nucleus | 0.17 | 1/d | fit |
| $k_{10}$ | YAP degradation rate | 1.8 | 1/d | fit |
| $k_{11}$ | pYAP degradation rate | 30 | 1/d | fit |
| $K_{tot}$ | Sum of amounts KIN, pKIN | 5.5 | $1/\mu m^3$ | free choice |

**Supplemental Experimental Procedures**

**3D reconstruction and spatial analysis of BC network diameter**

The BC network was reconstructed from 3D image stacks of CD13-stained tissue samples using the software MotionTracking as previously described (1,2). To reconstruct 3D image stacks of CD13, DAPI and phalloidin-stained tissue samples, a tile of 2 x 1 image stacks was stitched to cover an entire CV-PV axis. Then, the CD13 and DAPI channels were aligned to the 2-photon (DAPI + phalloidin) channel and image intensities were normalized as previously described (1). CD13 images were segmented using a local thresholding algorithm (maximum entropy), segmented objects were corrected for artefacts using standard morphological operations (opening/closing) and the triangulation mesh of the segmented surfaces was generated by the cube marching algorithm. A representation of the skeletonized image was generated using a 3D graph describing the geometrical and topological features of the bile canaliculi network. To reconstruct the CV and PV, the intensity of the DAPI and phalloidin channels were added and the vessels were segmented from the inverse signal.

For spatial analysis of the BC diameter, the CV-PV axis was computationally divided into 20 equidistant zones and canaliculi were assigned to the individual zones according to their relative position $\chi$ to the CV and PV using the following equation:

$$\chi = \frac{\mathrm{d}_{CV}}{\mathrm{d}_{CV} + \mathrm{d}_{PV}} * 20$$

Where $d_{CV}$ and $d_{PV}$ are the distance of BC from CV and PV, respectively. The BC diameter was quantified in the xy-plane and determined as average per zone. The zones directly adjacent to the CV and PV (zone 1 and 20, ~ 1 cell layer) were excluded from the analysis.

**Spatial analysis of nuclear YAP and PCNA and total YAP intensities within the CV-PV axis**

Nuclear YAP and PCNA as well as total YAP intensities were quantified from 2 x 1 image tiles covering an entire CV-PV axis. For nuclear quantifications, nuclei were segmented by DAPI intensity thresholding using the Fiji software and non-parenchymal cells were excluded based on size and circularity. Using the MotionTracking software, the lobule axis was computationally divided into 10 isocentric zones and the spatial position of the nuclei was calculated as described above (3D reconstruction and spatial analysis of BC network diameter). For total YAP quantifications, the average image intensity per zone was determined. For all quantifications, the median of the mean nuclear YAP and PCNA or total YAP intensities was calculated per zone. For each timepoint of a time course, the mean nuclear intensity for each zone from 3-5 CV-PV axes was quantified.

**Quantification of apical F-actin and pMLC density**

IF images of CD13 and F-actin or pMLC, acquired within the PV area, were segmented based on the apical marker CD13 using the mean shift (3) and the maximum entropy algorithms (4) implemented in the MotionTracking Software. To exclude BC lumens, intensity thresholding of the F-actin or pMLC intensity was additionally used. The mean F-actin or pMLC intensity was calculated per image and averaged for all images per time point. Within each condition (sham or PH), individual time courses were scaled to the mean value of all time courses by applying a scaling factor that was calculated as:

$$f_i = \frac{\sum_{j=1}^{N} y_{i,j} Y_j}{\sum_{j=1}^{N} Y_j^2}$$

Where $y_{i,j}$ is the intensity of the $i$-th curve in the $j$-th time point, $Y_j$ is the mean intensity of all curves in the $j$-th time point and $f_i$ is a scaling factor for the $i$-th curve, $Y_j = \frac{1}{K}\sum_{i=1}^{K} y_{i,j}$ , $N$ is the number of time points, $K$ is the number of the individual curves. The average intensity of all scaled time courses of one condition (sham or PH) was normalized to the untreated control (time point 0).

**Segmentation of BC and quantification of BC membrane length and perimeter from EM images**

To determine BC membrane length, BC membranes, including intra-luminal microvilli, were manually segmented from 10 x 10 EM image grids using the image analysis software Imod (http://bio3d.colorado.edu/imod). To estimate BC perimeter, the minimal enclosing ellipse of each BC segmentation was computationally determined as described previously (5). On average, 55 BC were quantified per sample.

**Quantification of the sub-apical and -basolateral gold particle density from immuno-EM images**

The apical and basolateral plasma membranes were segmented manually from 6 x 6 EM image grids (~ 30 x 30 µm) using the Imod software. Gold particles within the area 200 nm below the plasma membrane were quantified (see Fig. S5b) and gold particle density (particles per area) was calculated. For each EM image grid, the average gold particle density of all basolateral and apical membranes was determined.

**Mathematical model of YAP activation by cortical tension**

**Modeling strategy**

We considered a linear array of hepatocytes along the central-portal axis of a liver lobule. The individual hepatocytes were assumed to respond independently from each other to the local strength of a mechanical stimulus by YAP activation, i.e. increased concentration of YAP in the nucleus as compared to the untreated control. The mechanical stimulus was attributed to the osmotic pressure experienced by the apical membrane of hepatocytes that is proportional to the osmolyte concentration within the BC. We developed and coupled two sub-models to describe the changes of osmotic pressure and the concomitant activation of YAP after PH. Sub-model 1 is a biophysics-based model to predict the local mechanical stress that results from the alteration of osmolyte (bile acid) load in the BC network after PH. It considers the spatial geometry of the BC within the CV-PV axis of the lobule. Sub-model 2 is a biochemistry-based model that predicts the cellular response of YAP to the local mechanical stress. Model parameters are set as reported in the literature (1) or fitted to data reported here.

**Model definition**

**Sub-model 1 – Prediction of local mechanical stress at the apical domain of hepatocytes**

Bile flow through BC is driven by both peristaltic contractions of bile canaliculi and osmotic pressure, resulting from actively pumped bile salts and other osmolytes (2,6). Upon PH, with resection ratio r=62.8%±1.1% (mean and SEM) of liver mass without removal of the gall bladder, the total bile salt pool in the body is only marginally affected since intra-hepatic bile acids only account for 2-4% of the total bile acid pool (7). The body also retains the capacity of intestinal bile acid reabsorption. However, it increases the bile acid load in the liver remnant, which needs to transport the full bile salt pool through a proportionally reduced BC network. This increases the bile salt secretion flux per canaliculus. The concentration of osmolytes (*c*) within BC and consequently the osmotic pressure (*p*) have been shown to increase as the square root of the apical osmolyte secretion flux (2). Given that bile acids represent major osmolytes in bile (8), we here approximate *c* as the bile acid concentration in BC. The pressure magnitude is considered to be linearly dependent on the osmolyte concentration (9) while any spatial pressure dependencies are assumed as a common factor that cancels out in the ratio, see below. We predict the fluid pressure $p(x)$ within BC to increase relative to the pressure $p_0(x)$ of the control condition (sham operated mice) by the factor

$$\frac{p(x)}{p_0(x)} = \frac{c}{c_0} = \sqrt{\frac{100\%/(100\% - r)}{100\%/100\%}} = 1.64 \quad .$$

This increased intra-luminal pressure inflates the volume, and radius (a), of BC until it is counterbalanced by the Laplace pressure ($\tilde{p}$) exerted by the acto-myosin cortex. Assuming a cylindrical geometry of BC, the

Laplace pressure is given by $\tilde{p} = \tilde{\gamma}\ \frac{1}{a}$ where $\tilde{\gamma}$ is the surface tension or here cortical tension at the apical domain (for brevity termed cortical tension in the following). Since Laplace and fluid pressure are equal at steady state, we obtained a prediction for the *relative* increase of cortical tension, here denoted $\gamma(x)$, after PH as a function of position (*x*):

$$\gamma(x) = \frac{\tilde{\gamma}(x)}{\tilde{\gamma}_0(x)} = \frac{p(x)}{p_0(x)} * \frac{a(x)}{a_0(x)} = \sqrt{\frac{100\%}{100\% - r}} * \frac{a(x)}{a_0(x)} \qquad .$$

Note, the means of the measured values for *r*, *a*(*x*), $a_0(x)$ possess small SEM (below 10%) and we therefore approximate the SEM of the calculated cortical tension $\gamma(x)$ by propagating the SEM of the three measured quantities according to

$$SEM_{\gamma}(x) = \sqrt{\frac{100\%}{100\% - r}} * \frac{a(x)}{a_0(x)}$$

$$* \sqrt{\left(\frac{1}{2}\frac{100\%}{100\% - r}\right)^2 SEM_r{}^2(x) + \left(\frac{1}{a}\right)^2 SEM_a{}^2(x) + \left(\frac{1}{a_0}\right)^2 SEM_{a_0}{}^2(x)} \quad .$$

The predicted relative cortical tension levels of the bile canaliculi during regeneration are shown in Fig. S8a.

As a test of the predictive power of sub-model 1, we compared the predicted relative cortical tension to the measured relative pMLC levels (from Fig.2d) during regeneration. Since average pMLC levels were measured in the periportal 1/3 of the central-portal axis, they were compared to the averaged cortical tension values from the same spatial zones (Fig. S8b). Fig. S8b and a linear regression analysis reveal a high positive Pearson correlation (r=0.94) between the inferred cortical tension and pMLC levels, supporting sub-model 1. The diagonal line in Fig. S8b represents the linear curve fit $\frac{pMLC}{pMLC_0} = 1 + 0.62 *$

$(\gamma - 1)$. The predicted cortical tension was used as input to sub-model 2 to predict the local response of YAP.

**Sub-model 2 – Prediction of YAP activation by cortical tension**

The model of YAP activation by cortical tension is derived as follows and sketched in Fig. 7a. Solid (dashed) arrows in Fig. 7a denote reactions or transport steps (regulatory interactions), respectively. Depending on the position within the CV-PV axis, hepatocytes are exposed to different biliary pressures (2) and thus experience spatially heterogeneous cortical tension at the apical domain. We assumed that hepatocytes adapt to this spatially heterogeneous background cortical tension profile $\tilde{\gamma}_0(x)$ under normal conditions. Further, we assumed that a relative increase of cortical tension activates YAP through the Hippo pathway. In the liver, as in other tissues and species, YAP nuclear localization is regulated via phosphorylation by kinases of the Hippo signaling pathway (10–12). YAP phosphorylation by kinases, here termed KIN, causes its cytoplasmic retention (by binding to cytoplasmic sequestration factors, here termed SF) or degradation, thus preventing its translocation into the nucleus. Based on this, the model considers five YAP regulatory mechanisms: YAP synthesis, degradation, (de)phosphorylation, cytoplasmic retention and nuclear-cytoplasmic shuttling. Further, we use $\gamma(x)$, the ratio of the local cortical tension to the local background level of tension, as the mechanical stimulus that induces dephosphorylation of YAP kinases.

Altogether, sub-model 2 comprises 2 sub-cellular compartments (cytoplasm and nucleus), 11 reactions as numbered in Fig. 7a and 6 variables for species concentrations that depend on the spatial position within the CV-PV axis and time after PH. Reactions 1 and 2 are modelled according to Michaelis-Menten rate laws, accounting for the limited amounts of upstream kinases and phosphatases (parameters $k_1$, $K_{M1}$ and $k_2$, $K_{M2}$, respectively). The mechanical stimulus $\gamma(x)$ is a factor of the maximum dephosphorylation rate $k_1$. YAP synthesis is modeled as influx $k_3$. Within the lobule axis, hepatocytes are heterogenous with respect to their ploidy and nuclei number (1) as well as metabolic profile (13). This may affect cellular YAP protein levels. To account for potential spatial differences in YAP protein synthesis, we quantified the total YAP intensity profile at time point $t$=2d post PH as a proxy for relative (dimensionless) protein synthesis rate *s(x)* and inserted this experimentally measured spatial profile into the model equation for $k_3(x)=k_{3,0}*s(x)$.

All other reactions throughout the cytoplasm (numbers i=4-11) are modelled with simple mass action kinetics, each with a single rate constant $k_i$, see Table S2. The rate of the phosphorylation reaction 4 (Fig. 7a) was set proportional to the concentration of activated kinases [pKIN]. For transport between subcellular compartments of different volumes by reactions 8 and 9, scaling factors of volume ratios are derived from conservation of mass and introduced in the equations for cytoplasmic YAP ([YAP]) and nuclear YAP ([nYAP]).

The dynamics of the reaction network shown in Fig. 7a is modeled by 6 ordinary differential equations (ODE) for the 6 species' concentrations, as follows:

$$\frac{d[KIN]}{dt} = \frac{k_1 \cdot \gamma(x) \cdot [pKIN]}{K_{M1} + [pKIN]} - \frac{k_2 \cdot [KIN]}{K_{M2} + [\mathrm{KIN}]}$$

$$\frac{d[pKIN]}{dt} = -\frac{k_1 \cdot \gamma(x) \cdot [pKIN]}{K_{M1} + [pKIN]} + \frac{k_2 \cdot [KIN]}{K_{M2} + [\mathrm{KIN}]}$$

$$\frac{d[YAP]}{dt} = k_{3,0} \cdot s(x) - k_4 \cdot [pKIN] \cdot [YAP] + k_5 [pYAP] + k_8 \cdot \frac{V_N}{V_C} \cdot [\mathrm{nYAP}] - k_9 \cdot [YAP] - k_{10} \cdot [YAP]$$

$$\frac{d[pYAP]}{dt} = k_4 \cdot [pKIN] \cdot [YAP] - \mathrm{k}_5 [pYAP] - k_6 [pYAP] + k_7 [pYAP_{SF}] - k_{11} \cdot [\mathrm{p}YAP]$$

$$\frac{d[pYAP_{SF}]}{dt} = k_6 \cdot [pYAP] - k_7 [pYAP_{SF}]$$

$$\frac{d[nYAP]}{dt} = k_9 \cdot \frac{V_C}{V_N} \cdot [\mathrm{YAP}] - k_8 [nYAP]$$

$$[\mathrm{YAP}_{total}] = \frac{V_C}{V_C + V_N} \cdot \left([YAP] + [pYAP] + [pYAP_{SF}]\right) + \frac{V_N}{V_C + V_N}[nYAP]$$

The last, algebraic equation provides the average total concentration of all YAP forms as an observable that can be compared to measured average YAP intensity data.

For model analysis, we considered two time scales, one fast (minutes to hours) of protein modifications and turnover versus one slow (days) time scale of tissue growth. On the fast time scale, model simulations always converged to a unique stable steady state for a given mechanical stimulus. Assuming that the mechanical stimulus changes on the slow time scale, the state of the model adapted accordingly, rendering it a quasi steady state. To keep the model simple, we considered temporally constant and spatially uniform parameter values.

**Parameter estimation**

The model comprises 16 parameters. Of these, the compartment volumes of nuclei and cytoplasm, have previously been measured (1) and are set to the published values as referenced in Table S2. As revealed by the analytical model analysis, five of these parameters affect the steady state solution only in parameter ratios of forward and backward rates of the four reversible reactions, hence individually these parameters are non-identifiable from given steady state data. We therefore set the value of five selected parameters to 1, see Table S2. The remaining 9 parameters were estimated by fitting the quasi steady state solution of the model to 120 independent data points from two observables on 6 experimental conditions (t=0d, t=0.8d, 1.5d, 2.0d, 3.0d, 5.0d) at 10 different spatial locations along the central-portal axis. We used a combination of Evolutionary Programming and Levenberg-Marquardt as global and local optimizers as implemented in the software Copasi (14). Parameter optimization was performed in the software Copasi until convergence and results were confirmed by simulations in the software Morpheus (15).

**Model results**

Analyzing the 6 ordinary differential equations at steady state, we predicted nuclear YAP levels as a function of cortical tension $\gamma(x,t)$. We found a unique solution that represents a stable steady state:

$$[nYAP](\gamma) = \frac{V_C}{V_N} * \frac{k_9}{k_8} * \frac{k_3}{k_{10}} * \frac{1}{1 + \frac{2k_2 K_{M1} * {}^{k_4}/_{k_{10}}}{\left(1 + {}^{k_5}/_{k_{11}}\right) * f(\gamma)}}$$

$$[YAP_{total}](\gamma) = \frac{V_C}{V_C + V_N} * \frac{k_3}{k_{11}} * \left(1 + \frac{k_6}{k_7}\right) + \frac{V_C}{V_C + V_N} * \frac{k_3}{k_{10}} * \frac{1 + \frac{k_9}{k_8} - \frac{k_{10}}{k_{11}} * \left(1 + \frac{k_6}{k_7}\right)}{1 + \frac{2k_2 K_{M1} * {}^{k_4}/_{k_{10}}}{\left(1 + {}^{k_5}/_{k_{11}}\right) * f(\gamma)}}$$

$$f(\gamma) = \gamma k_1 \left(1 + \frac{K_{M2}}{K_{tot}}\right) + k_2 \left(\frac{K_{M1}}{K_{tot}} - 1\right) + \sqrt{\left(\gamma k_1 \left(1 + \frac{K_{M2}}{K_{tot}}\right) + k_2 \left(\frac{K_{M1}}{K_{tot}} - 1\right)\right)^2 - 4(\gamma k_1 - k_2) k_2 \frac{K_{M1}}{K_{tot}}}$$

This closed analytical form of the steady state solution allowed us to understand the role of individual parameters in the response of hepatocytes and guided the parameter estimation strategy. Specifically, given steady state data for nuclear and total YAP levels, only a subset of the parameters is individually identifiable while others only affect the solution through ratios of parameters and were chosen for computational convenience (see Table S2).

Employing this spatially and temporally constant parameter set (Table S2) for all 120 available data points, the model predicted nuclear (see Fig. S8c) and total (see Fig. S8d) YAP profiles from position- and time-dependent cortical tension $\gamma(x,t)$ and the position-dependent but temporally constant proxy for protein synthesis (see Fig. S8d, red curve). Overall, we observe a very good match between the confidence intervals of data and model prediction, both indicated by SEM intervals in Figs. S8c and S8D, where the calculated SEM intervals of $\gamma(x,t)$ (see Fig. S8a) have been propagated to SEM intervals for YAP. Spatial fluctuations in the model prediction are ultimately a response to spatial fluctuations in the experimentally measured radius profiles that are used to predict $\gamma(x,t)$. Just for the last time point, t=5d, the model prediction overestimates for both nuclear and total YAP levels.

Having established the mechanistic model of YAP turnover, activation and subcellular translocation, we next asked how sensitive the YAP response is with respect to the level of cortical tension. To study this question, we inserted a spatially averaged protein synthesis rate (s=<*s(x)*>=1.5) and considered $\gamma$ as a control parameter. The resulting stimulus-response curve (see Fig. 7b) reveals a plateau around the normal unstressed condition $\gamma < 1.8$ followed by a sigmoidal part with a slope of approximately 500% and a plateau at high stimulus levels. Note, the scatter around the theoretical dose-response curve is attributable to the spatial variation in protein synthesis rate that was not considered in the spatially averaged model prediction for s=<*s(x)*>.

Such a sigmoidal stimulus-response curve of nuclear YAP levels ensures that normal (e.g. diet-triggered) fluctuations in biliary fluid pressure and hence cortical tension are tolerated and do not mount a proliferative response while a stronger pressure increase robustly activates proliferation in a switch-like manner.

**Statistics**

For quantification of BC diameter (Fig. 1b, 6b), BC perimeter and membrane length (Fig. 1d, 1e), apical F-actin and pMLC density (Fig. 2b, 2d) as well as nuclear YAP and PCNA intensity (Fig. 3c, 3d, 6d), *n* represents the number of mice per condition or timepoint. For quantification of sub-apical and -basolateral YAP density from EM images (Fig. S5c), *n* represents the number of EM image grids analyzed. For quantification of nuclear and cytoplasmic intensity of YAP from IF images of hepatocyte cultures (Fig. 5b, 5e, S7a) as well as pMLC levels from Western blot (Fig. S7b, c), *n* represents the number of experiments. Measurements are given as mean +/- standard error of the mean (s.e.m.; Fig. 1b, 2b, 2d, 3c, 3d, 5b, 5e, 6b, 6d, S7, S8) or box-whisker plot with median, 25-75 quartiles and minimum/maximum error bars (Fig. 1d, 1e, S5c). The significance of the difference of two conditions was estimated by Student's t-test. To calculate the significance of the difference of two spatial or temporal profiles a and b (Fig. 1b, 2b, 2d, 3c, 3d, 6b, 6d), the normalized differences $\Delta_i$ of two values $a_i$ and $b_i$ was calculated using

$$\Delta_i = \frac{(a_i - b_i)}{\sqrt{{\sigma_{a_i}}^2 + {\sigma_{b_i}}^2}}$$

where *i* is the zone index within the CV-PV axis or time point of a time course. The Student's t-test was applied assuming as null hypothesis that the mean difference is 0.

**Software**

The MotionTracking software (1) (http://motiontracking.mpi-cbg.de) was used for analysis and 3D reconstructions of IF images as well as statistical analysis. Imod (http://bio3d.colorado.edu/imod) was used for manual segmentation of EM images. GraphPad (GraphPad Software, Inc.) was used for graphical representations. Fiji (16) was used for background subtraction of IF images, image visualization and Western blot quantifications. Copasi (14) was used for parameter estimation in the mathematical sub-model 2. Morpheus (15) was used to simulate and analyze the coupled mathematical models.
